# NG2 glia protect against prion neurotoxicity by inhibiting prostaglandin E2 signaling

**DOI:** 10.1101/2023.04.04.535590

**Authors:** Yingjun Liu, Jingjing Guo, Maja Matoga, Marina Korotkova, Per-Johan Jakobsson, Adriano Aguzzi

**Affiliations:** Institute of Neuropathology, University of Zurich, Switzerland; Department of Medicine, Karolinska Institutet, Sweden

**Author notes:** Correspondence Adriano Aguzzi or Yingjun Liu: Institute of Neuropathology, University of Zurich, Schmelzbergstrasse 12, 8091 Zurich, Switzerland. or.

## Abstract

Oligodendrocyte-lineage cells, including NG2 glia, undergo prominent changes in various neurodegenerative disorders. This raises the question of how myelinating cells interact with neurodegenerative processes. Here, we found that NG2 glia were activated after prion infection in cerebellar organotypic cultured slices (COCS) and in brains of prion-inoculated mice. In both model systems, depletion of NG2 glia exacerbated prion-induced neurodegeneration and accelerated prion pathology. Loss of NG2 glia unleashed a microglial reaction promoting the biosynthesis of prostaglandin E2 (PGE2), which augmented prion neurotoxicity in the HovS cell line, primary neurons and COCS through binding to the EP4 receptor. Single-cell RNA sequencing revealed molecular signatures of inflammatory, disease-associated and MHC^+^ microglia but not of interferon-responsiveness in PGE2-producing microglia of prion-inoculated mice. Pharmacological or genetic inhibition of PGE2 biosynthesis attenuated prion-induced neurodegeneration in COCS and mice, reduced the enhanced neurodegeneration in NG2-glia-depleted COCS after prion infection, and dampened the acceleration of prion disease in NG2-glia-depleted mice. These data unveil a non-cell-autonomous interaction between NG2 glia and microglia in prion disease and suggest that PGE2 signaling may represent an actionable target against prion diseases.

## Introduction

Neurodegenerative diseases such as Alzheimer’s disease (AD), Parkinson’s disease (PD) and prion diseases involve multiple cell types and various genetic and environmental factors (Wareham, Liddelow et al. 2022). Accumulating evidence suggest that both cell-autonomous and non-cell-autonomous mechanisms contribute to the neurodegenerative process (Aguzzi and Liu 2017, Liu and Aguzzi 2019). Among the common neurodegenerative disorders, prion diseases can be best mimicked with laboratory animal models (Brandner and Jaunmuktane 2017). Prion-inoculated mice develop a fatal neurodegenerative condition that is almost indistinguishable from its human counterpart at the neuropathological level, providing an invaluable model system for investigating the cellular and molecular mechanisms of chronic neurodegeneration relevant to humans.

Different neurodegenerative diseases have distinct initial pathogenic triggers and clinical manifestations, yet certain molecular and cellular alterations, such as abnormal protein aggregation and activation of glial cells, are common to all these disorders (Wareham, Liddelow et al. 2022). Large amounts of data support vital roles of microglia and astrocytes in the initiation and progression of neurodegeneration. However, the function of other cell types, including oligodendrocyte-lineage cells, in the pathogenesis of neurodegenerative diseases has been elusive, and investigations on this crucial aspect were largely ignored by the neurodegenerative disease community. Recent advances in high resolution gene expression analyses have enabled a deeper understanding of this complex cellular landscape. In addition to confirming the alterations of microglia and astrocytes, single-cell and spatial transcriptomics analyses of brain tissues from patients and animal models have unraveled prominent changes of oligodendrocyte linage cells in multiple neurodegenerative disorders (Mathys, Davila-Velderrain et al. 2019, Chen, Lu et al. 2020, Pandey, Shen et al. 2022), including prion diseases (Dimitriadis, Zhang et al. 2022, Slota, Sajesh et al. 2022).

Oligodendrocyte-lineage cells are traditionally considered as supportive cells in the central nervous system (CNS), producing myelin sheaths that insulate nerve fibers and help speed up transmission of electrical signals along neuronal axons. It is unclear how myelinating cells interact with other cell types of the brain in the context of neurodegenerative process. Here we investigated the role of oligodendrocyte precursor cells (NG2 glia) in chronic neurodegeneration induced by prion infections. We found that NG2 glia were neuroprotective and played a crucial role in influencing the microglial pathway that is responsible for the biosynthesis of prostaglandin E2 (PGE2), which promotes prion-induced neurodegeneration through binding to the EP4 receptor. These data suggest that NG2 glia have an impact on an intricate cell-cell interaction network in prion disease, and highlight NG2 glia and PGE2 signaling as potential targets for disease-modifying therapies against neurodegenerative disorders.

## Results

### NG2 glia activation in prion disease models

Our previous gene expression analyses of prion-inoculated mice (Sorce, Nuvolone et al. 2020) and prion-infected cerebellar organotypic cultured slices (COCS) (Liu, Senatore et al. 2022), as well as recent single-cell transcriptomics from other groups (Dimitriadis, Zhang et al. 2022, Slota, Sajesh et al. 2022), point to possible transcriptional changes in NG2 glia during prion disease. However, it is unclear how these cells respond to prion infections at the cellular level. We therefore examined the markers of NG2 glia, including NG2 and platelet-derived growth factor receptor alpha (Pdgfrα), in prion-infected COCS and mouse brains by western blotting and immunofluorescence. We found an increase in NG2 protein levels in both paradigms (**Fig. 1a-d**). The protein level of Pdgfrα was also upregulated in mouse brains after prion infection, although not as much as NG2 (**Fig. 1c-d**). Similarly, we observed enhanced NG2 immunoreactivity in prion-infected C57BL/6J and Tga20 COCS as well as in prion-inoculated mouse brains (**Fig. 1e-h** and **Supplementary Fig. 1a-b**). After prion infection, NG2 glia exhibited enlargement of cell bodies and arborization of cellular processes (**Fig. 1g**), reminiscent of the NG2 glia phenotypes in mouse models of brain injuries (Alonso 2005, Jin, Riew et al. 2018), suggesting that neuronal damage plays a role in prion-induced NG2 glia activation.

**Figure 1.**
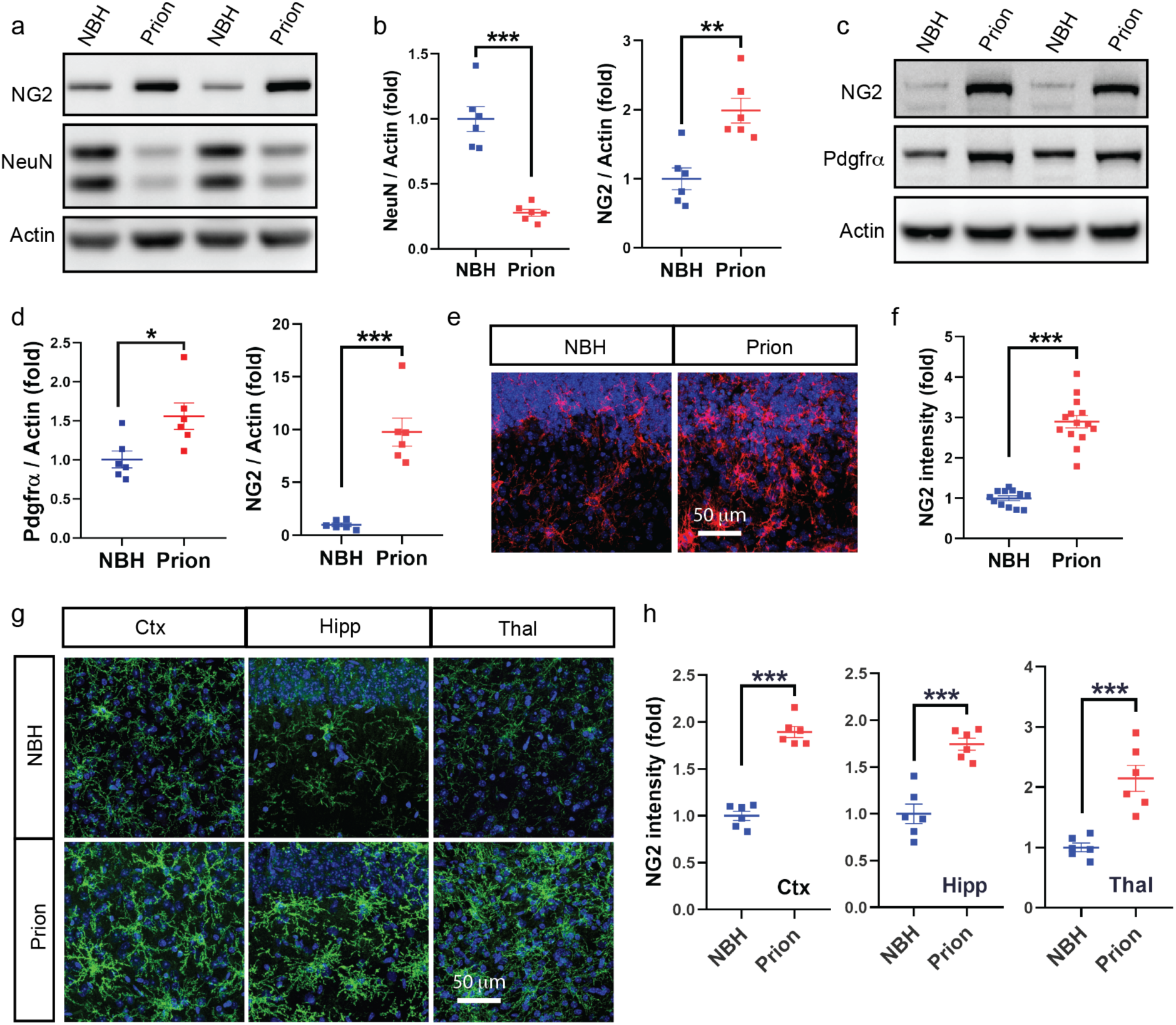
NG2 glia activation in prion disease models. **a**-**b**, Western blots (**a**) and quantification (**b**) of NG2 and NeuN in Tga20 COCS exposed to prions or non-infectious brain homogenates (NBH, here and in subsequent figures). n = 6 samples/condition; 6-8 slices/sample. **c**-**d**, Western blots (**c**) and quantification (**d**) of NG2 and Pdgfrα in brain tissues of mice inoculated with prions or NBH. n = 6 mice/condition. **e**, NG2 immunofluorescence showing NG2 glia activation in prion-infected Tga20 COCS vs Tga20 COCS exposed to NBH. Nuclei were stained with DAPI (blue), here and henceforth. **f**, Quantification of NG2 immunointensity shown in **e**. n = 12 slices for NBH; n = 14 slices for Prion. **g**, NG2 glia activation in the cerebral cortex (Ctx), hippocampus (Hip) and thalamus (Thal) of prion-inoculated mice vs mice inoculated with NBH. **h**, Quantification of NG2 immunointensity shown in **g**. n = 6 mice/group. Here and in following figures: *: p < 0.05; **: p < 0.01; ***: p < 0.001; n.s: not significant.

### Loss of NG2 glia enhances prion neurotoxicity and accelerates prion disease

To facilitate functional investigations of NG2 glia, we previously developed efficient and selective NG2 glia depletion strategies ex vivo and in vivo (Liu and Aguzzi 2020). We used the PDGFR inhibitor CP673451 to deplete NG2 glia from COCS, and combined cell-type selective diphtheria toxin receptor (DTR) expression with systemic DT injections for in vivo NG2 glia depletion (Liu and Aguzzi 2020).

To determine the functional relevance of NG2 glia in prion-induced neurodegeneration, we depleted NG2 glia from COCS by continuous (**Supplementary Fig. 2a-b**) or transient (**Supplementary Fig. 3a**) CP673451 treatment in the presence or absence of prions, and compared them to DMSO-treated controls. Again, we saw enhanced NG2 immunoreactivity in prion-infected C57BL/6J and Tga20 COCS and – as expected – NG2 glia loss in both uninfected and prion-infected COCS after continuous CP673451 treatment (**Fig. 2a-b** and **Supplementary Fig. 2c-d**). Although NG2 glia depletion did not influence neuronal survival in the absence of prions, it enhanced neurodegeneration in prion-exposed C57BL/6J and Tga20 COCS (**Fig. 2a-b** and **Supplementary Fig. 2c-d**), suggesting a neuroprotective role of NG2 glia under pathological conditions. A similar enhancement of neurodegeneration was also observed in prion-infected COCS after transient CP673451 treatment (**Supplementary Fig. 3b-c**). Interestingly, NG2 glia number was largely restored 3 weeks after transient CP673451 treatment in prion-infected COCS (**Supplementary Fig. 3b-c**), suggesting that short-term destruction of NG2 glia suffices to influence prion neurotoxicity.

**Figure 2.**
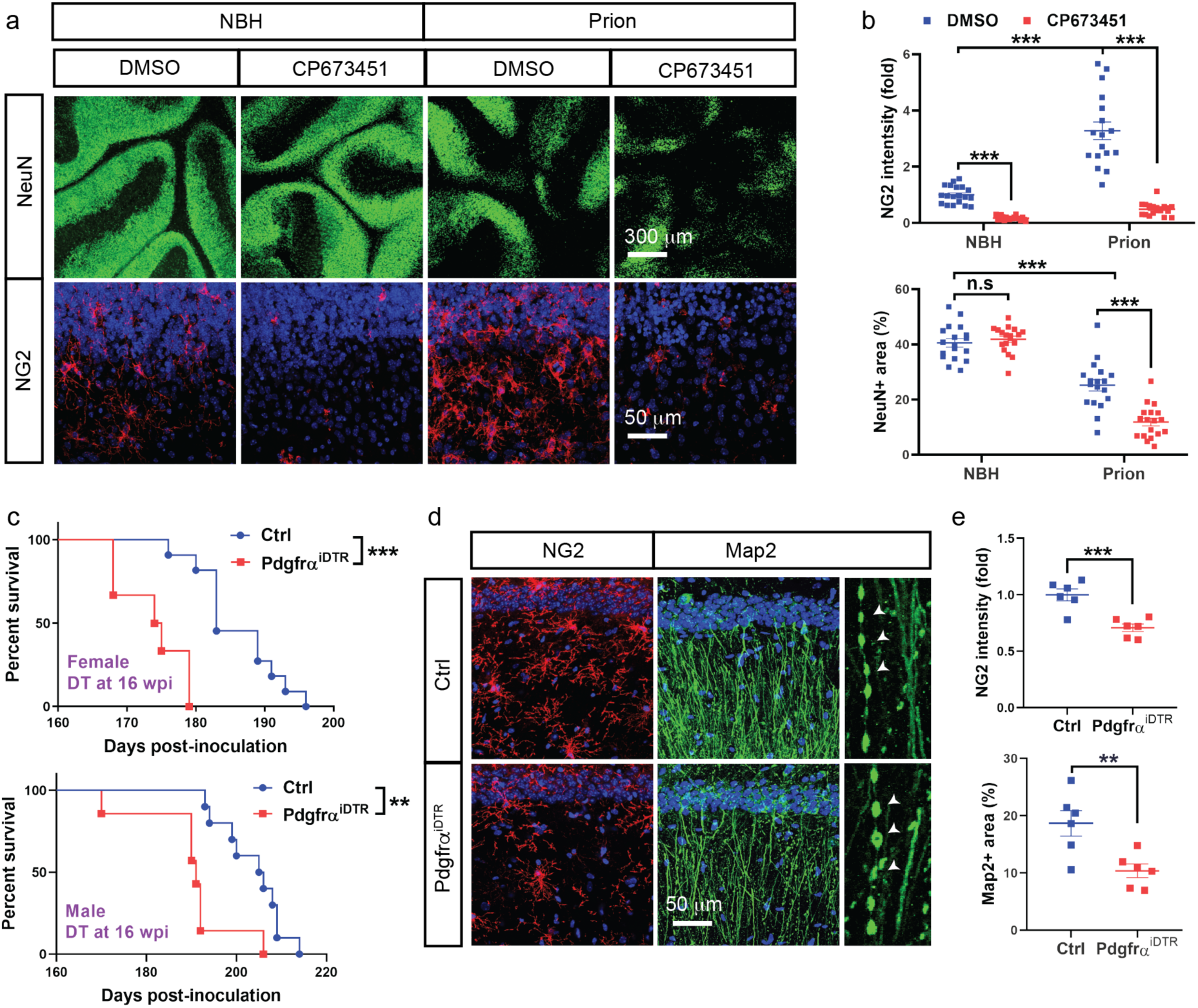
NG2 glia depletion enhances prion neurotoxicity and accelerates prion disease. **a**, NeuN and NG2 immunofluorescence showing prion-induced neurodegeneration in NG2-glia-depleted (CP673451) and NG2 glia intact (DMSO) Tga20 COCS. **b**, Quantification of NG2 immunointensity and NeuN positive area shown in **a**. n > 16 slices/condition. **c**, Survival curves showing accelerated prion disease in NG2-glia-depleted (Pdgfrα^iDTR^) mice after prion inoculation. Median survival: 191 days for male Pdgfrα^iDTR^ mice; 205.5 days for male control mice; 174.5 days for female Pdgfrα^iDTR^ mice; 184 days for female control mice. NG2 glia depletion was induced at 16 wpi. **d**, NG2 and Map2 immunofluorescence showing enhanced dendritic pathology in hippocampi of NG2-glia-depleted (Pdgfrα^iDTR^) mice after prion inoculation. Arrowheads: pathologic dendrites with varicosities and fragmentation. NG2 glia depletion was induced at 16 wpi; brain samples were collected at 21 wpi. **e**, Quantification of NG2 immunointensity and Map2 positive area shown in **d**. n = 6 mice/group.

To investigate the effects of NG2 glia depletion on prion disease in vivo, we crossed Pdgfrα-CreER mice with iDTR mice and generated Pdgfrα-CreER/iDTR (Pdgfrα^iDTR^) mice. Pdgfrα-CreER and iDTR littermates were used for controls. We inoculated mice with prions one month after tamoxifen treatment, and administered DT daily for 5 days at 12 or 16-weeks post inoculation (wpi) (**Supplementary fig. 4a**). According to immunofluorescent examinations, there was ∼50% decrease of NG2 glia number in the brains of Pdgfrα^iDTR^ mice compared to controls after DT injection (Liu and Aguzzi 2020). Prion disease was accelerated in NG2-glia-depleted Pdgfrα^iDTR^ mice of both sexes when NG2 glia depletion was induced at 16 wpi (**Fig. 2c**), and in male mice when NG2 glia depletion was induced at 12 wpi (**Supplementary Fig. 4b**), possibly attributable to the sex effects on the incubation time of prion disease in animal models (Loeuillet, Boelle et al. 2010, Akhtar, Wenborn et al. 2011).

Early neurodegeneration in prion disease is characterized by severe dendritic pathology (Fuhrmann, Mitteregger et al. 2007), which can be best visualized in the hippocampus. To investigate whether loss of NG2 glia affects prion-induced neurodegeneration in vivo, we inoculated a second cohort of Pdgfrα^iDTR^ and control mice after tamoxifen treatment, administered DT at 16 wpi and collected brains 4 weeks thereafter at ∼21 wpi. Immunostaining of NG2 confirmed reduction of NG2 glia number in Pdgfrα^iDTR^ mice compared to controls (**Fig. 2d-e**). We then examined hippocampal dendritic pathology by Map2 immunofluorescence. As expected, we observed dendritic damages characterized by dendritic varicosities and fragmentation in both NG2 glia intact and NG2-glia-depleted mice after prion infection (**Fig. 2d**). However, the density of surviving dendrites was less in Pdgfrα^iDTR^ mice than in control mice (**Fig. 2d-e**), suggesting that NG2 glia deficiency also enhances prion-induced neurodegeneration in vivo. HE staining indicated similar spongiform changes in the brains of prion-inoculated Pdgfrα^iDTR^ mice and control mice (**Supplementary Fig. 5**).

### Loss of NG2 glia does not influence PrP^C^ expression, prion replication and prion-induced neuroinflammation

PrP^C^ is the substrate of prion replication (Bueler, Aguzzi et al. 1993), and is essential for prion disease pathogenesis (Brandner, Isenmann et al. 1996). Tissue abundance of PrP^C^ correlates with the incubation time of prion disease in animal models (Fischer, Rulicke et al. 1996, Minikel, Zhao et al. 2020). We therefore examined the levels of PrP^C^ and PrP^Sc^ (the protease K-resistant pathological PrP) in NG2-glia-depleted COCS and mice by western blotting. We found that neither the PrP^C^ (**Supplementary Fig. 6a-b**) nor the PrP^Sc^ levels (**Supplementary Fig. 6c-d**) noticeably changed in COCS treated with CP673451 compared to controls. In addition, similar levels of PrP^C^ (**Supplementary Fig. 6e-f**) and PrP^Sc^ (**Supplementary Fig. 6g-h**) were observed in the brains of NG2-glia-depleted Pdgfrα^iDTR^ mice and control mice. Hence, enhanced neurodegeneration and accelerated prion disease after NG2 glia depletion were not caused by alterations of PrP^C^ expression and prion replication.

We previously found that NG2 glia were required for maintaining the homeostatic microglia state (Liu and Aguzzi 2020). However, depletion of NG2 glia in cultured brain slices and adult mice did not influence microglia numbers or induce notable changes of tissue-level neuroinflammatory responses under normal conditions (Liu and Aguzzi 2020). To determine whether loss of NG2 glia affects prion-induced microglia activation and neuroinflammation, we compared the density of Cd68^+^ reactive microglia between NG2-glia-depleted and intact COCS in the presence or absence of prions. We found that reactive microglia were rare in control COCS, but their number increased after prion infection (**Supplementary Fig. 7a-b**). Nevertheless, the levels of microglia activation were comparable between NG2-glia-depleted and intact COCS, with or without prion infection (**Supplementary Fig. 7a-b**). Similarly, we did not observe any changes of Iba1 and Cd68 immunoreactivity in NG2-glia-depleted brains after prion inoculation (**Supplementary Fig. 7c-d**). Furthermore, the mRNA levels of the proinflammatory factors, tumor necrosis factor alpha (TNFα), Interleukin 1 beta (IL1β) and IL12β, were unaltered by NG2 glia depletion in COCS and mouse brains after prion infection (**Supplementary Fig. 8a-b**). Collectively, these data suggest that loss of NG2 glia does not change prion-induced microglia activation and neuroinflammation at tissue level.

### Loss of NG2 glia enhances the pathway that is responsible for PGE2 biosynthesis

When evaluating gene expression changes in NG2-glia-depleted COCS with RNA sequencing (Liu and Aguzzi 2020), we noticed possible dysregulation of genes related to the biosynthesis of PGE2, which was reported to be increased in the cerebral spinal fluid (CSF) of prion disease patients (Minghetti, Greco et al. 2000, Minghetti, Cardone et al. 2002).

To confirm the potential influence of NG2 glia on PGE2, we depleted NG2 glia in COCS with CP673451, and examined the expression levels of cyclooxygenase 2 (Cox2) and prostaglandin E synthase (Ptges), two major enzymes responsible for PGE2 biosynthesis under pathological conditions. Both immunofluorescence and qRT-PCR results confirmed efficient NG2 glia depletion after CP673451 treatment (**Fig. 3a-c**). Importantly, we found that the expression levels of both Cox2 and Ptges were upregulated in NG2-glia-depleted COCS compared to NG2 glia intact COCS (**Fig. 3c**). Furthermore, we treated COCS with PDGFAA, the ligand for Pdgfrα, which promotes NG2 glia proliferation (Calver, Hall et al. 1998). Immunofluorescent and qRT-PCR results indicated increase of NG2 glia density in PDGFAA-treated COCS compared to controls (**Fig. 3d-f**). We found that increase of NG2 glia number decreased the expression levels of both Cox2 and Ptges (**Fig. 3f**). These data indicate that NG2 glia regulate the PGE2 biosynthesis pathway. In line with the above findings, upregulation of both Cox2 and Ptges were also observed in NG2-glia-depleted COCS after prion infection (**Fig. 3g**). As comparison, we found that neither Cox1 nor Ptgds changed upon NG2 glia depletion (**Supplementary Fig. 9**).

**Figure 3.**
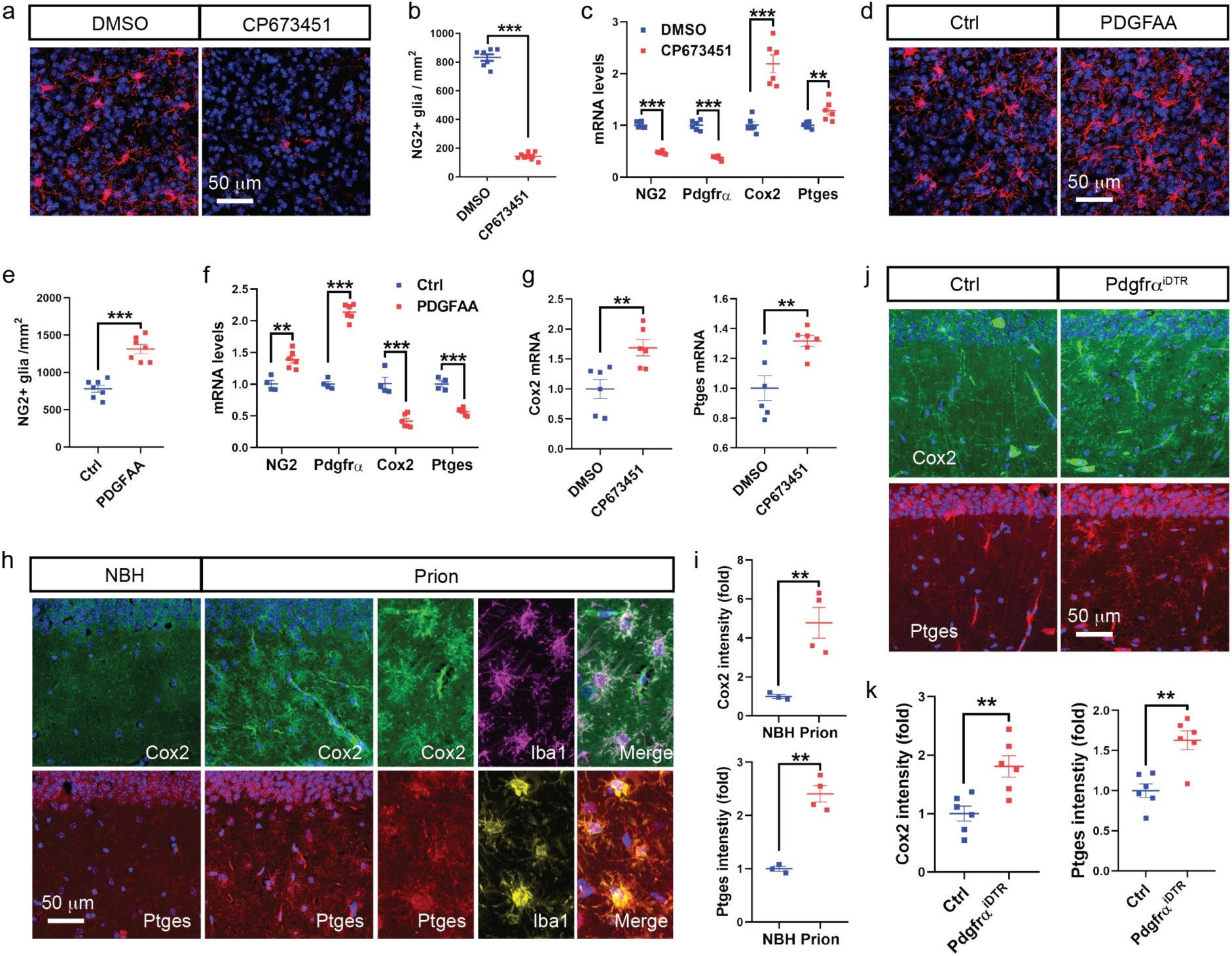
Loss of NG2 glia upregulates Cox2-Ptges expression. **a**-**b**, NG2 immunofluorescence (**a**) and quantification (**b**) in intact (DMSO) and NG2-glia-depleted (CP673451) C57BL/6J COCS. n = 7 slices/condition. **c**, qRT-PCR results showing downregulation of NG2 and Pdgfrα and upregulation of Cox2 and Ptges in NG2-glia-depleted (CP673451) C57BL/6J COCS. n = 6 samples; 6-8 slices/sample. **d**-**e**, NG2 immunofluorescence (**d**) and quantification (**e**) in control (Ctrl) and PDGFAA-treated C57BL/6J COCS. n = 7 slices/condition. **f**, qRT-PCR results showing upregulation of NG2 and Pdgfrα and downregulation of Cox2 and Ptges in C57BL/6J COCS with increased NG2 glia density (PDGFAA) compared to NG2 glia intact (Ctrl) C57BL/6J COCS. n = 6 samples; 6-8 slices/sample. **g**, qRT-PCR results showing upregulation of Cox2 and Ptges in NG2-glia-depleted (CP673451) Tga20 COCS after prion exposure. n = 6 samples; 6-8 slices/sample. **h**, Cox2, Ptges and Iba1 immunofluorescence in the hippocampus showing upregulation of Cox2 and Ptges and their colocalization with microglia in brains of terminally sick prion-inoculated mice. **i**, Quantification of Cox2 and Ptges immunointensity shown in **h**. n = 3 mice for NBH and n = 4 mice for prion. **j**, Cox2 and Ptges immunofluorescence in the hippocampus showing upregulation of Cox2 and Ptges in the brains of NG2-glia-depleted (Pdgfrα^iDTR^) mice compared to NG2 glia intact (Ctrl) mice after prion inoculation. **k**, Quantification of Cox2 and Ptges immunointensity shown in **j**. n = 6 mice/group.

To investigate whether loss of NG2 glia induces similar changes of Cox2 and Ptges in vivo, we examined the expression levels of Cox2 and Ptges as well as their cellular localization in the brain by immunofluorescence. We found that the immunoreactivity of both Cox2 and Ptges were relatively low under normal condition (**Fig. 3h-i**); however, prion infection increased their expression levels (**Fig. 3h-i**), especially in microglia (**Fig. 3h**). Furthermore, NG2 glia depletion increased prion-induced upregulation of Cox2 and Ptges (**Fig. 3j-k**), suggesting that NG2 glia also regulate this pathway in vivo.

### NG2 glia regulate microglial PGE2 pathway through multiple mechanisms

To molecularly characterize PGE2-producing microglia in prion-infected brains, we analyzed a previously generated single-cell RNA sequencing dataset (Slota, Sajesh et al. 2022) based on a prion infection model similar to that used in the current study. In normal brains, microglia were largely (∼99%) Cox2^-^ or Ptges^-^; however, the fractions of Cox2^+^ and Ptges^+^ microglia were increased after prion infection in both the cerebral cortex (**Fig. 4a-b**) and hippocampus (**Supplementary Fig. 10a-b**). Expression analysis identified 943 differentially expressed genes (DEGs) between Cox2^+^ and Cox2^-^ microglia (**Supplementary table 1**), and 847 DEGs between Ptges^+^ and Ptges^-^ microglia **(Supplementary table 2**), respectively, in the cerebral cortex of prion-infected mice (**Fig. 4c**). The majority (∼80%-90%) of DEGs identified in Cox2^+^ microglia were also dysregulated in Ptges^+^ microglia (**Fig. 4c**), suggesting that Cox2^+^ and Ptges^+^ microglia largely overlap. Similar results were also observed in the hippocampus of prion-infected mice (**Supplementary Fig. 10c, Supplementary table 3** and **Supplementary table 4**).

**Figure 4.**
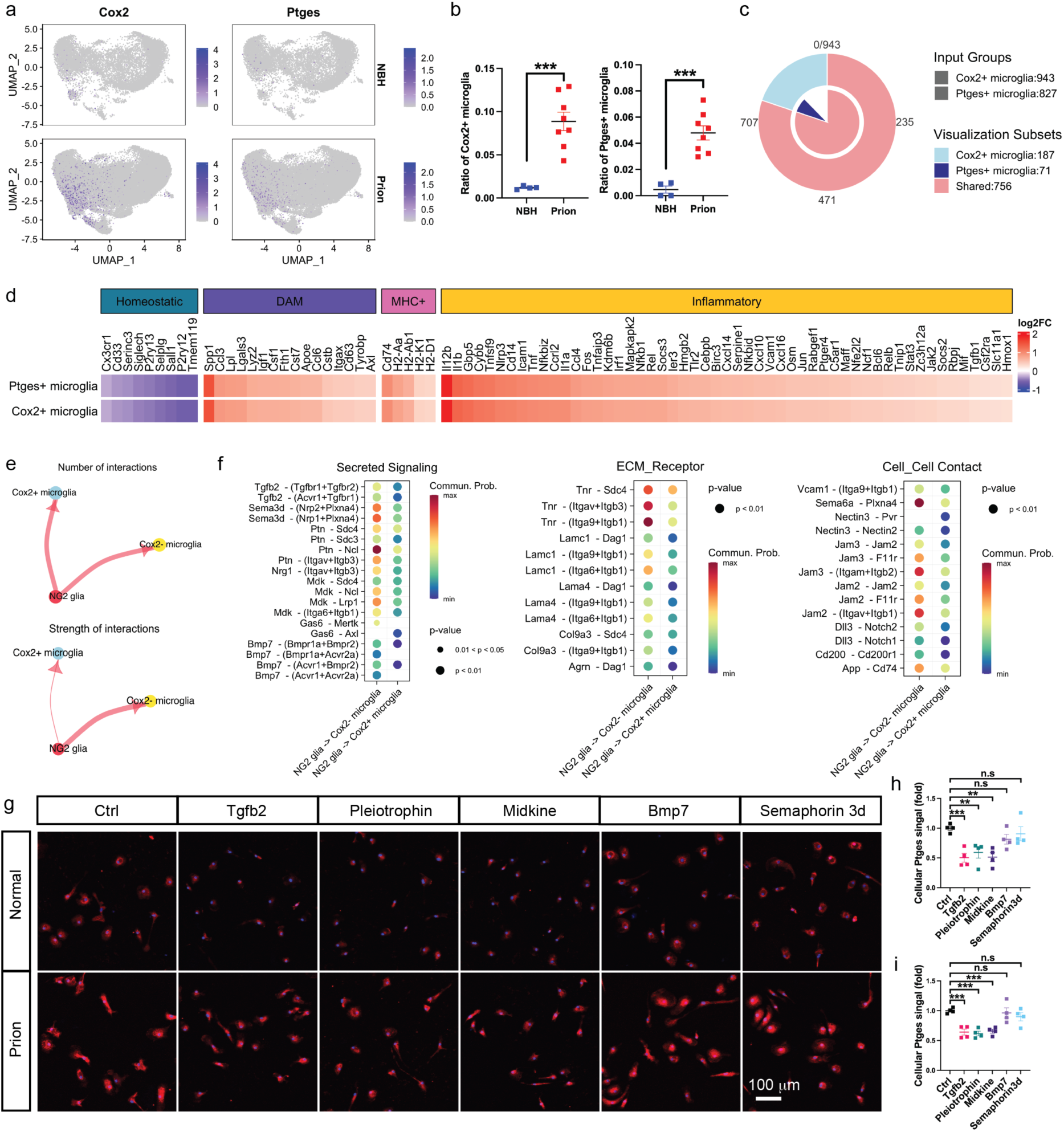
NG2 glia regulate microglial Cox2-Ptges through multiple mechanisms. **a**, UMAP of single-cell RNA-seq data showing Cox2^+^ and Ptges^+^ microglia among total microglia in the cerebral cortex of prion- or NBH-inoculated mice. **b**, Quantification of Cox2^+^ and Ptges^+^ microglia fractions against total microglia in the cerebral cortex of prion- or NBH-inoculated mice shown in **a**. n = 4 mice for NBH and n = 8 mice for Prion. **c**, Venn diagram showing numbers of shared and distinct DEGs of Cox2^+^ and Ptges^+^ microglia in the cerebral cortex of prioninoculated mice. **d**, Heatmap showing downregulation of homeostatic microglia signature genes and upregulation of DAM and MHC^+^ microglia signature genes as well as inflammatory genes in Cox2^+^ and Ptges^+^ microglia in the cerebral cortex of prion-inoculated mice. **e**, Cell-Chat analysis of cell-cell communications showing unaltered number of interactions (plotted as the thickness of the edges) but reduced strength of interactions (plotted as the thickness of the edges) from NG2 glia to Cox2^+^ microglia in the cerebral cortex of prion-inoculated mice. **f**, Heatmaps showing significantly weakened NG2 glia to Cox2^+^ microglia interaction pathways in the cerebral cortex of prion-inoculated mice. **g**, Immunofluorescence showing that NG2-glia-derived factors such as Tgfb2, Pleiotrophin and Midkine but not Bmp7 and Semaphorin3d suppress Ptges expression in primary microglia in the presence or absence of prions. **h**, Quantification of Ptges immunointensity in microglia in the absence of prions shown in **g**. n = 4 independent experiments. **i**, Quantification of Ptges immunointensity in microglia in the presence of prions shown in **g**. n = 4 independent experiments.

The signature genes of homeostatic microglia, including Tmem119, P2ry12 and Cx3cr1, were downregulated in Cox2^+^/Ptges^+^ microglia (**Fig. 4d** and **Supplementary Fig. 10d**), whereas disease-associated microglia (DAM) (Keren-Shaul, Spinrad et al. 2017) signature genes such as Itgax, Cst7 and Apoe as well as MHC^+^ microglia (Mathys, Adaikkan et al. 2017) signature genes such as H2-D1, H2-Aa and Cd74 were upregulated in Cox2^+^/Ptges^+^ microglia (**Fig. 4d** and **Supplementary Fig. 10d**). No interferon response microglia (Mathys, Adaikkan et al. 2017) signature genes were significantly altered in Cox2^+^/Ptges^+^ microglia (**Fig. 4d**). In addition, we found that Cox2^+^/Ptges^+^ microglia expressed higher levels of inflammatory genes (**Fig. 4d** and **Supplementary Fig. 10d**). These data suggest that Cox2^+^/Ptges^+^ microglia represent either a novel microglia population or a mixture of different microglia populations that are involved in neurodegenerative diseases.

To investigate the mechanisms by which NG2 glia influence the generation of Cox2^+^/Ptges^+^ microglia in prion disease, we analyzed the cell-cell communication between NG2 glia and Cox2^+^ or Cox2^-^ microglia at the single-cell level with CellChat (Jin, Guerrero-Juarez et al. 2021). We found that the number of interactions between NG2 glia and Cox2^+^ or Cox2^-^ microglia was unchanged (**Fig. 4e** and **Supplementary Fig. 10d**). However, the strength of interactions between NG2 glia and Cox2^+^ microglia was weakened compared to NG2 glia and Cox2^-^ microglia (**Fig. 4e**). The weakened NG2 glia inputs on Cox2^+^ microglia were associated with multiple mechanistic categories, including secreted signaling such as transforming growth factor beta 2 (Tgfb2), pleiotrophin (Ptn) and midkine (Mdk), ECM-receptor interaction such as tenascin R (Tnr), laminin subunit gamma 1 (Lamc1) and collagen type IX alpha 3 chain (Col9a3), and cell-cell contact such as vascular cell adhesion protein 1 (Vcam1), nectin cell adhesion molecule 3 (Nectin3) and junctional adhesion molecule 3 (Jam3) (**Fig. 4f** and **Supplementary Fig. 10f**).

We previously reported that Tgfb signaling disruption in NG2-glia-depleted COCS may be responsible for the loss of homeostatic microglia state (Liu and Aguzzi 2020). These observations not only validate the Cellchat analysis results (**Fig. 4f** and **Supplementary Fig. 10f**), but also suggest that some of the weakened NG2 glia to Cox2^+^ microglia signaling might be causally linked to the enhanced cellular state transition from Cox2^-^ microglia into Cox2^+^ microglia in the NG2-glia-depleted mouse brains after prion infection. To test this hypothesis, we established high-purity serum-free microglia cultures from mouse brains through Cd11b immunopanning (**Supplementary Fig. 11a-b**). After three days in culture, we treated primary microglia with several of the NG2-glia-derived factors identified by the cell-cell communication analysis (**Fig. 4f** and **Supplementary Fig. 10f**), including Tgfb2, Ptn, Mdk, bone morphogenetic protein 7 (Bmp7) and semaphorin3d (Sema3d) in the presence or absence of prions, and examined Ptges levels with immunofluorescence. We found that Tgfb2, Ptn, Mdk but not Bmp7 and Sema3d suppressed microglial Ptges expression under both experimental conditions (**Fig. 4g-i**). These data suggest that NG2 glia can directly influence microglial Cox2-Ptges pathway through multiple mechanisms.

### Inhibition of PGE2 biosynthesis diminishes prion neurotoxicity and decelerates prion disease

Although previous studies indicated that the level of PGE2 in the CSF was increased in prion disease patients (Minghetti, Greco et al. 2000, Minghetti, Cardone et al. 2002), it is unclear whether PGE2 plays a causal role in prion-induced neurodegeneration. To investigate this, we treated normal and prion-infected COCS with PGE2, PGD2, or DMSO as control, and examined neurodegeneration by NeuN immunofluorescence. PGE2 treatment had no effect on neuronal survival under normal conditions but enhanced neuronal death in prion-infected COCS (**Fig. 5a-b**). In contrast, prion-induced neurodegeneration in PGD2-treated COCS was largely unaltered compared to DMSO (**Fig. 5a-b**). To investigate whether inhibition of PGE2 biosynthesis protects against prion-induced neurodegeneration, we treated normal and prion-infected COCS with C118 or C934 (Larsson, Steinmetz et al. 2019), two high-selective Ptges inhibitors. Both Ptges inhibitors reduced neurodegeneration in prion-exposed COCS (**Fig. 5c-d**). These results suggest that PGE2 is a potent enhancer of prion neurotoxicity, and inhibition of PGE2 biosynthesis is neuroprotective.

**Figure 5.**
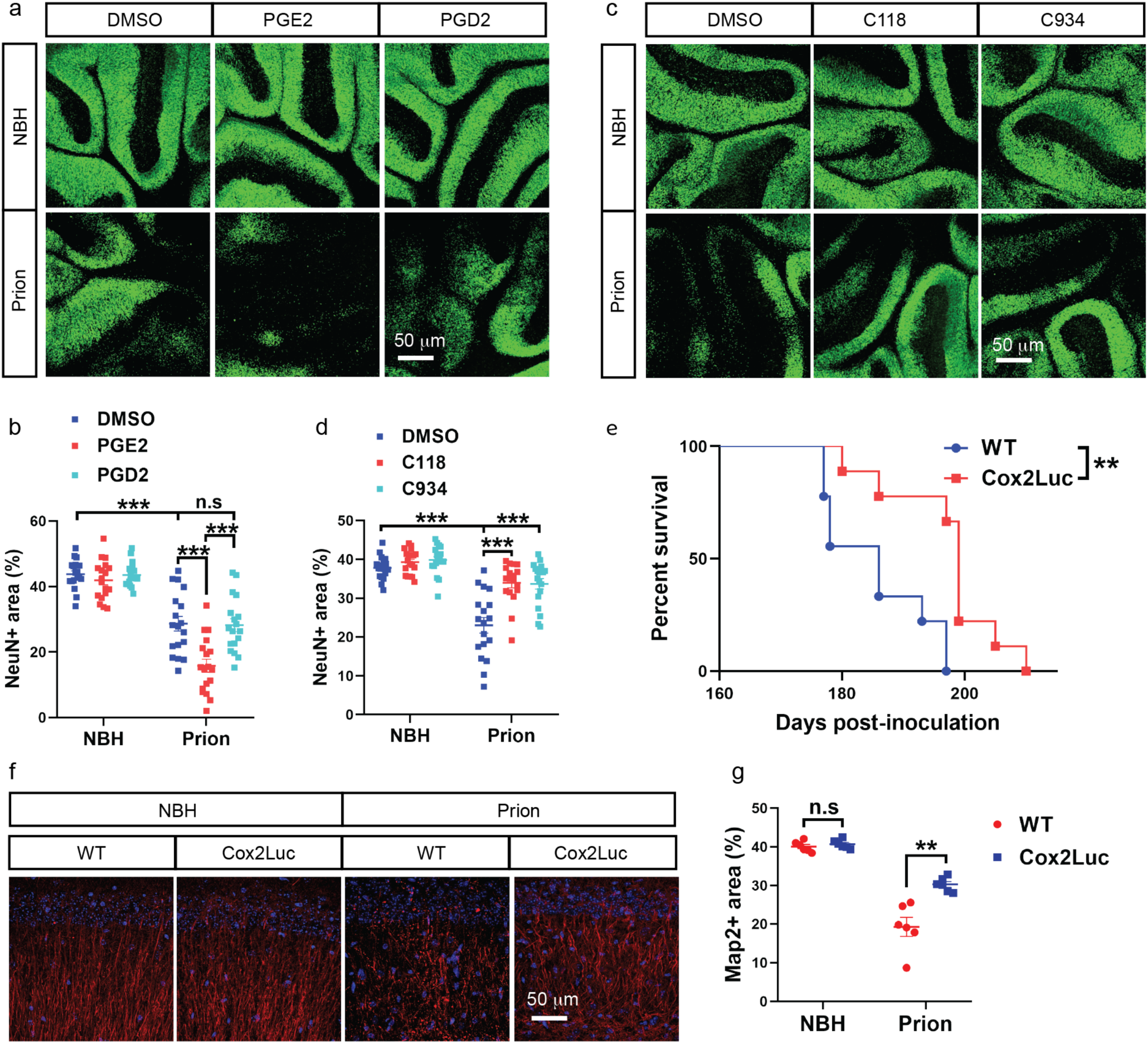
Cox2-Ptges inhibition diminishes prion neurotoxicity and decelerates prion disease. **a**-**b**, NeuN immunofluorescence (**a**) and quantification (**b**) showing enhanced neurodegeneration in PGE2-treated compared to DMSO-treated and PGD2-treated Tga20 COCS after prion infection. n = 18 slices/condition. **c**-**d**, NeuN immunofluorescence (**c**) and quantification (**d**) showing diminished neurodegeneration in Tga20 COCS treated with Ptges inhibitors C118 and C934 compared to DMSO-treated Tga20 COCS after prion infection. n = 18 slices/condition. **e**, Survival curves showing decelerated prion disease in Cox2 knockout (Cox2Luc) mice compared to littermate WT mice. Median survival: 186 days for control mice; 199 days for Cox2Luc mice. **f**-**g**, Map2 immunofluorescence (**f**) and quantification (**g**) showing diminished dendritic pathology in the hippocampi of Cox2Luc mice after prion inoculation compared to littermate WT mice. Brain samples were collected at 21 wpi. n = 6 mice/group.

To investigate the effects of PGE2 biosynthesis inhibition on prion disease in vivo, we inoculated Cox2Luc mice, in which the Cox2 gene was replaced with the coding sequence of luciferase, and control mice with prions. We found that Cox2Luc mice survived longer than their littermate controls after prion infection (**Fig. 5e**), suggesting that elevation of Cox2 and PGE2 production during prion disease are detrimental in vivo. In consistent with this, immunofluorescent staining of Map2 demonstrated ameliorated prion-induced dendritic pathology in the hippocampus of the Cox2Luc mice compared to controls (**Fig. 5f-g).**

### Inhibition of PGE2 biosynthesis rescues the enhanced neurodegeneration and accelerated prion disease after NG2 glia depletion

Next, we investigated whether the upregulation of Cox2 and Ptges and the subsequent increase of PGE2 biosynthesis were responsible for the enhanced prion neurotoxicity and accelerated prion disease after NG2 glia depletion. We examined neuronal survival upon inhibition of Ptges activity by C118 and C934 in normal and prion-infected COCS with or without NG2 glia depletion. Consistent with the aforementioned findings (**Fig. 2a-b**), loss of NG2 glia enhanced prion-induced neurodegeneration (**Fig. 6a-b**). However, this enhancement was largely blocked by C118 and C934 treatment (**Fig. 6a-b**). These data suggest that the enhanced neurodegeneration in NG2-glia-depleted COCS is mediated by PGE2.

**Figure 6.**
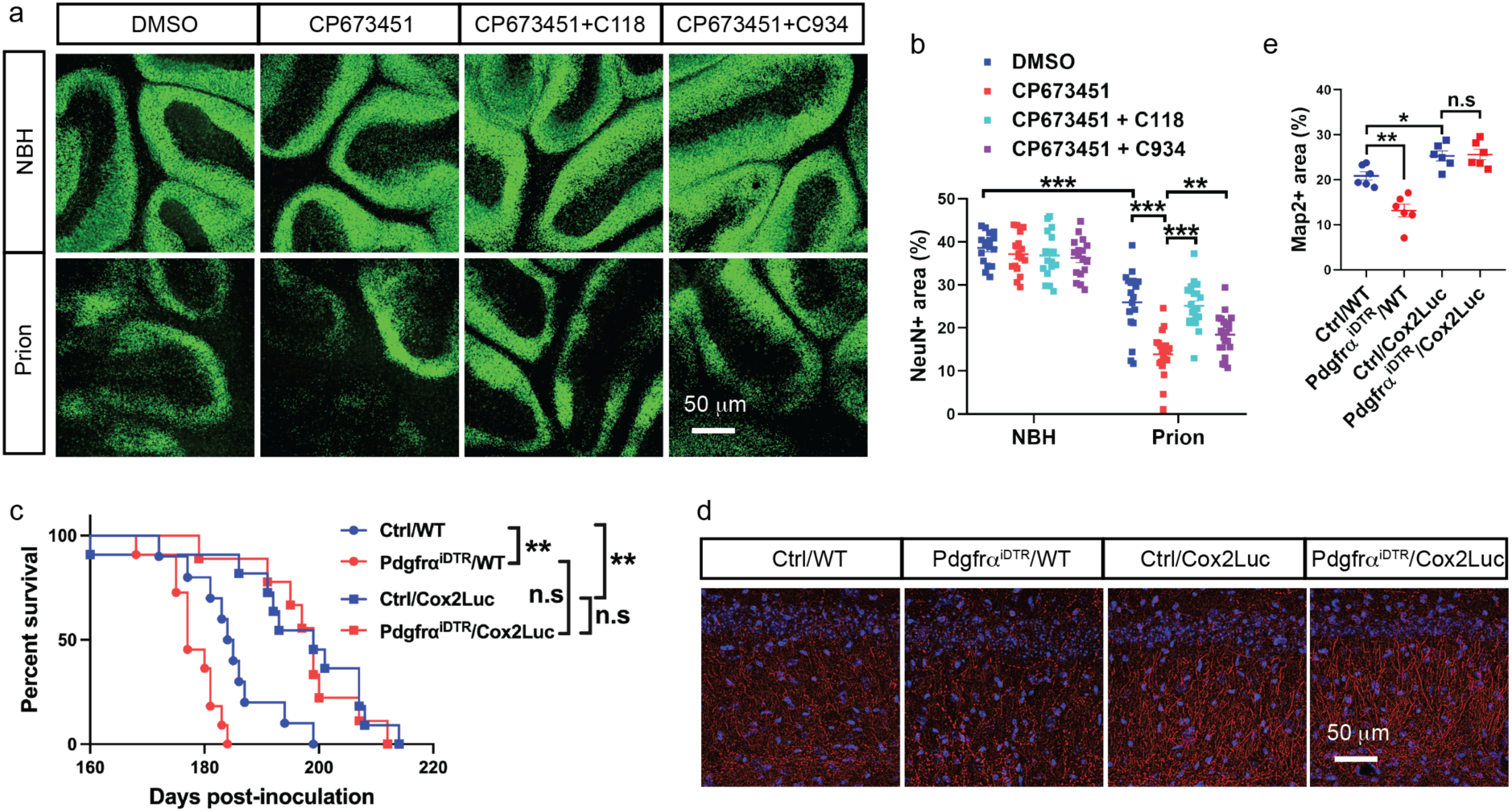
Cox2-Ptges inhibition rescues enhanced neurodegeneration and accelerated prion disease after NG2 glia depletion. **a-b**, NeuN immunofluorescence (**a**) and quantification (**b**) showing that enhanced neurodegeneration in prion-infected, NG2-glia-depleted (CP673451) Tga20 COCS can be rescued by treatment with Ptges inhibitors C118 and C934. n = 16 slices/condition. **c**, Cox2 ablation (Cox2Luc) suppresses the acceleration of prion disease in NG2-glia-depleted (Pdgfrα^iDTR^) mice. Median survival: 177 days for Pdgfrα^iDTR^/WT mice; 184.5 days for Ctrl/WT mice; 199 days for Pdgfrα^iDTR^/Cox2Luc and Ctrl/Cox2Luc mice. NG2 glia depletion was induced at 16 wpi. **d-e**, Map2 immunofluorescence (**d**) and quantification (**e**) showing enhanced dendritic pathology in NG2-glia-depleted (Pdgfrα^iDTR^) hippocampi of prion-infected mice, and its rescue by Cox2 ablation (Cox2Luc). n = 6 mice/group.

We then crossed Pdgfrα-CreER mice and iDTR mice with Cox2Luc mice to generate Pdg-frα^iDTR^/Cox2Luc mice. Pdgfrα-CreER/Cox2Luc and iDTR/Cox2Luc mice served as controls (Ctrl/Cox2Luc). We infected Pdgfrα^iDTR^/Cox2Luc and Ctrl/Cox2Luc mice with prions and compared them to prion inoculated Pdgfrα^iDTR^ and genetically matched control mice (designated as Pdgfrα^iDTR^/WT and Ctrl/WT mice, respectively). All mice were treated with tamoxifen one month before prion inoculation and injected with DT at 16 wpi. As expected, loss of NG2 glia accelerated prion disease in Pdgfrα^iDTR^/WT mice compared to Ctrl/WT mice (**Fig. 6c**). However, the accelerated prion disease in NG2-glia-depleted mice was almost completely rescued in Pdgfrα^iDTR^/Cox2Luc mice compared to Ctrl/Cox2Luc mice (**Fig. 6c**), and both Pdg-frα^iDTR^/Cox2Luc mice and Ctrl/Cox2Luc mice survived longer than Pdgfrα^iDTR^/WT and Ctrl/WT mice (**Fig. 6c**). Loss of Cox2 also rescued the enhanced prion-induced dendritic pathology in Pdgfrα^iDTR^/Cox2Luc mice compared to Pdgfrα^iDTR^/WT mice (**Fig. 6d-e**). These findings confirm that increased PGE2 biosynthesis underlies the enhanced prion neurotoxicity and accelerated prion disease in NG2 glia deficient mice.

### PGE2 enhances prion neurotoxicity through the EP4 receptor

To investigate how PGE2 promotes prion-induced neurodegeneration, we first examined the cellular localization of its receptors, including prostaglandin E receptor 1 (Ptger1, also known as EP1), Ptger2 (also known as EP2), Ptger3 (also known as EP3) and Ptger4 (also known as EP4), in the adult mouse brain by immunofluorescence. We found that all four PGE2 receptors were expressed by neurons (**Supplementary Fig. 12**). To determine which receptor might be responsible for the enhanced prion neurotoxicity, we expressed them one by one together with GFP in HovS cells chronically infected with prions (Avar, Heinzer et al. 2020) through lentiviral transduction, and examined death of HovS cells in the presence or absence of PGE2. We found that the number of GFP^+^ HovS cells were comparable between PGE2- and DMSO-treated groups when Ptger2, Ptger3 or a control construct was expressed (**Fig. 7a-b**). However, expression of Ptger1 and Ptger4 reduced the survival of HovS cells in PGE2- treated groups compared to DMSO-treated controls (**Fig. 7a-b**). Furthermore, we observed an abnormal morphology of surviving HovS cells, characterized by retraction of cellular processes and shrinkage of cell bodies, after PGE2 treatment only when Ptger1 and Ptger4 were expressed (**Fig. 7a**). Therefore, PGE2 may enhance prion-induced neurodegeneration through activating Ptger1- and Ptger4-mediated signaling.

**Figure 7.**
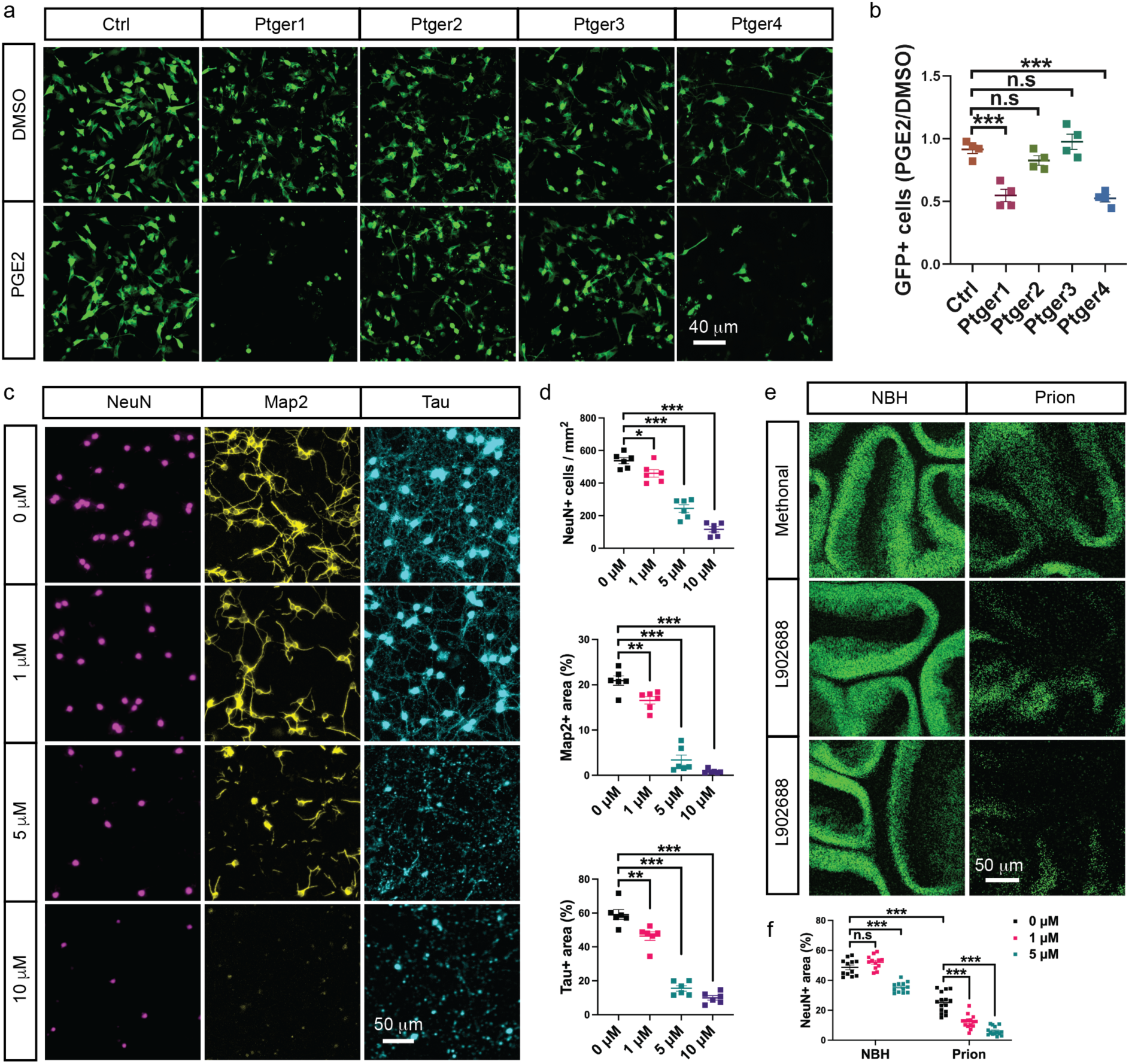
PGE2 enhances prion neurotoxicity mainly through the EP4 receptor (Ptger4)**. a-b**, Live-cell imaging (**a**) and quantitative analysis (**b**) of chronically prion-infected HovS cells expressing control (Ctrl) transgene or one of the four PGE2 receptors (Ptger1-4). Effects of PGE2 treatment on prion-induced cell toxicity were measured with the ratio of GFP signals under the PGE2 condition against the DMSO condition. n = 4 independent experiments; 4 technical repeats/experiment. **c**, Immunofluorescence of NeuN, Map2 and Tau showing cellular damages of prion-infected primary neurons treated with different concentrations of Ptger4 agonist L902688. **d**, Quantification of neuronal density as well as Map2 positive and Tau positive areas shown in **c**. n = 6 independent experiments. **e-f**, NeuN immunofluorescence (**e**) and quantification (**f**) showing concentration-dependent enhancement of prion-induced neurodegeneration in L902688-treated Tga20 COCS. n = 12 slices/condition.

To further confirm the possible involvement of Ptger1 and Ptger4 in PGE2-mediated enhancement of neurodegeneration after prion infection, we established primary neuronal cultures using cerebellum from C57BL/6J mice. Immunofluorescence indicated expression of both Ptger1 and Ptger4 in primary neurons (**Supplementary Fig. 13**). We infected primary neurons with prion-containing brain homogenates at day 5, and treated them with different concentrations of Ptger1 agonist 17-phenyl-trinor-PGE2 (17-pt-PGE2), Ptger4 agonist L902688 or control solvents 10 days later for 48 hours. We found that L902688 enhanced the death of prion-infected neurons and damages to neuronal processes in a concentration dependent manner (**Fig. 7c-d**). High concentration of L902688 also induced neuronal death in the absence of prions (**Supplementary Fig. 14a-b**), suggesting activation of Ptger4 signaling is highly toxic to neurons. In contrast, we found that activation of Ptger1 with several concentrations of 17-pt-PGE2 had no effects on neuronal survival in the presence of prions (**Supplementary Fig. 14c-d**). Similarly, we found that activation of Ptger4 but not Ptger1 enhanced prion-induced neurodegeneration in COCS in a concentration-dependent manner (**Fig. 7e-f** and **Supplementary Fig. 15a-b**). These results confirm that PGE2 enhances prion-induced neurodegeneration primarily by activating the Ptger4-mediated signaling.

## Discussion

Oligodendrocyte lineage cells have long been considered as supportive components of the CNS and bystanders in neurogenerative diseases (Ettle, Schlachetzki et al. 2016). However, accumulating evidence suggest that this cell lineage may have additional functions other than producing myelin sheath (Funfschilling, Supplie et al. 2012, Liu and Zhou 2013, Fruhbeis, Kuo-Elsner et al. 2020), and may be actively involved in neurodegenerative processes (Lee, Morrison et al. 2012, Pandey, Shen et al. 2022). Here we found that loss of oligodendrocyte precursor cells unleashed a microglial pathway that is responsible for PGE2 biosynthesis, which enhances prion-induced neurotoxicity and accelerates prion disease in animal models (**Supplementary Fig. 16**). These data uncover a crucial role of oligodendrocyte precursor cells in a non-cell-autonomous interaction network during prion infection, and support a protective function of these cells during chronic neurodegeneration. Developing approaches that promote the beneficial activities of oligodendrocyte precursor cells may hold potential for novel disease-modifying therapies.

The study of functions of oligodendrocyte-lineage cells was limited by the lack of efficient and specific ways to manipulate them ex vivo and in vivo. To overcome this difficulty, we previously established experimental approaches to deplete oligodendrocyte precursor cells in cultured brain slices and in adult mouse brains (Liu and Aguzzi 2020), which formed the basis for the discoveries described here. Adaption of these tools to other models of neurodegeneration, such as AD and PD, would facilitate the investigations of disease-specific roles of oligodendrocyte precursor cells.

We found that depletion of oligodendrocyte precursor cells augmented the PGE2 biosynthesis pathway in cultured brain slices and adult mouse brains after prion infection, revealing a previously unknown regulatory mechanism of PGE2 signaling in the CNS. The role of PGE2 signaling in prion disease and prion-induced neurodegeneration was undefined, but PGE2 levels were found to be increased in the CSF of prion disease patients (Minghetti, Greco et al. 2000, Minghetti, Cardone et al. 2002). Our findings indicate that PGE2 signaling is a strong enhancer of prion neurotoxicity and a potent driver of prion disease development, which underlies the enhanced neurodegeneration and accelerated prion disease in mice deficient of oligodendrocyte precursor cells. Furthermore, we found that PGE2 enhanced prion-induced neurodegeneration through directly binding to its receptor EP4 on neuronal cells. These data suggest that Ptges inhibitors blocking PGE2 biosynthesis, such as C118 and C934 (Larsson, Steinmetz et al. 2019), which were tested in the current study, as well as EP4 signaling antagonists, may represent good drug candidates for the treatment of prion disease.

Microglia play important roles in prion disease pathogenesis (Zhu, Herrmann et al. 2016, Carroll, Race et al. 2018, Bradford, McGuire et al. 2022). Besides neurons, glial cells, especially activated microglia, also express PGE2 receptors (Bonfill-Teixidor, Otxoa-de-Amezaga et al. 2017). Manipulation of PGE2 signaling in microglia influences their reaction in models of neurodegeneration (Johansson, Pradhan et al. 2013, Woodling, Wang et al. 2014). Therefore, in addition to directly acting on neurons, PGE2 may modulate microglial phenotypes through autocrine signaling, and contribute to neurodegeneration indirectly.

We previously reported that oligodendrocyte precursor cells were required for maintaining the homeostatic state of microglia in the adult mouse brain; however, their loss did not result in microglia activation and neuroinflammation under normal conditions (Liu and Aguzzi 2020), in contrast to NG2 glia depletion in rats (Nakano, Tamura et al. 2017). While we now confirm this even in the context of prion infections, we found that deficiency of oligodendrocyte precursor cells dysregulates the PGE2 biosynthesis pathway in microglia. Single-cell RNA sequencing data indicate that PGE2-producing microglia exhibit the molecular features of previously reported DAM (Keren-Shaul, Spinrad et al. 2017) and of MHC^+^ microglia (Mathys, Adaikkan et al. 2017) but not interferon responsive microglia (Mathys, Adaikkan et al. 2017), and express higher levels of inflammatory genes. PGE2-producing microglia may represent a novel microglia population of diseased brains, or a mixture of several different microglia populations identified previously. Profiling larger numbers of microglial cells in the prion-inoculated brain with single-cell RNA sequencing in future studies may help clarify this crucial question.

Using CellChat (Jin, Guerrero-Juarez et al. 2021) to analyze NG2 glia-microglia communications in the prion-infected brain at the single-cell level, we identified multiple weakened NG2-glia-to-microglia signals that might be mechanistically linked to the enhanced generation of PGE2-producing microglia in NG2-glia-depleted brains after prion infection. Of these, we validated Tgfb2, Ptn and Mdk experimentally in primary microglia cultures. Targeting these signals or other CellChat-identified NG2-glia-derived factors may represent promising ways for modulating microglia phenotypes in neurodegenerative diseases.

## Materials and methods

### Mouse experiments

C57BL/6J mice were obtained from Charles River, Germany. Tga20 mice (Fischer, Rulicke et al. 1996) were obtained from Laboratory Animal Services Center at University of Zurich, Switzerland. Pdgfrα-CreER mice (Stock No: 018280), iDTR mice (Stock No: 007900) and Cox2Luc mice (Stock No: 030853) were obtained from the Jackson Laboratory, USA. Double transgenic Pdgfrα-CreER/iDTR (Pdgfrα^iDTR^) mice were generated by crossing the Pdgfrα-CreER mice with the iDTR mice. To induce Cre activity and DTR expression in NG2 glia, Pdgfrα^iDTR^ mice were fed with tamoxifen-containing diet (ENVIGO, TD. 55125.I) for 4 weeks as previously reported (Liu and Aguzzi 2020). To generate Pdgfrα^iDTR^ mice on a Cox2Luc background, Pdg-frα-CreER mice and iDTR mice were first crossed with the Cox2Luc mice, then the resulting Pdgfrα-CreER mice and iDTR mice on the Cox2Luc background were crossed with each other. Littermates of the experimental groups were used as controls. All animal experiments in the current study were performed according to Swiss federal guidelines and had been approved by the Animal Experimentation Committee of the Canton of Zurich under the permits 040/2015, 139/2016, 243/2018 and 236/2019.

### Prion inoculation

Intracerebral prion inoculation was performed as previously described (Liu, Senatore et al. 2022). Briefly, adult mice were anesthetized with isoflurane and injected in the right hemisphere of the brain with 30 μl 0.01% w/v brain homogenates derived from adult C57BL/6J mice suffering from terminal prion disease. Prion strain used in the current study was the Rocky Mountain Laboratory strain of scrapie, passage 6. Mice inoculated with 30 μl 0.01% w/v non-infectious brain homogenates were used as controls. After prion inoculation, the health status of mice was closely monitored, and body weights of mice were recorded once per week. For experiments evaluating survival time, mice were euthanized when showing terminal disease symptoms or reached more than 20% loss of body weight. For biochemical and immunohistochemical analysis, mice were euthanized at specific time points after prion inoculation for tissue collection. To avoid potential influences of tamoxifen on prion disease development, mouse strains (e.g., Pdgfrα-CreER, iDTR and Pdgfrα^iDTR^ mice on WT or Cox2Luc background) went through tamoxifen food treatment was inoculated at least 4 weeks after switching back to the normal diet.

### Cerebellar organotypic cultured slices (COCS)

COCS were prepared according to a previously published protocol (Falsig and Aguzzi 2008). Briefly, cerebella from 12-day old C57BL/6J or Tga20 pups were dissected, embedded in low melting point agarose (Invitrogen, 15517-022) and cut into 350-μm thick slices with a vibratome (Leica, VT1000S) in cold Gey’s balanced salt solution (GBSS) supplemented with the glutamate receptor antagonist kynurenic acid (1 mM, Sigma, K3375) and glucose (33.33 mM, Sigma, G8769). Slices with intact morphology were collected and briefly washed 3 times in GBSS supplemented with kynurenic acid and glucose. Afterwards, brain slices were exposed to either 0.01% w/v prion-containing brain homogenates or 0.01% w/v brain homogenates derived from healthy mice for 1 hour at 4 °C and washed 5 times in GBSS supplemented with kynurenic acid and glucose. Six to eight slices were put on a Millicell-CM Biopore PTFE membrane insert (Millipore, PICM 03050) and kept on slice culture medium containing 50% MEM, 25% BME, 25% inactivated horse serum, 0.65% w/v glucose, 1% GlutaMax (ThermoFisher Scientific, 35050061) and 1% penicillin/streptomycin (ThermoFisher Scientific, 10378016) at 37 °C in a tissue culture incubator. Culture medium was changed 3 times per week.

### Treatment of brain slice cultures

Prostaglandin E2 (PGE2, sc-201225), prostaglandin D2 (PGD2, sc-201221) and EP1 receptor agonist 17-Phenyl-trinor-prostaglandin E2 (17-pt-PGE2, sc-201255) were purchased from Santa Cruz Biotechnology. EP4 receptor agonist L902688 (HY-119163) was purchased from MedChemExpress. Prostaglandin E synthase (Ptges) inhibitors C118 and C934 (Larsson, Steinmetz et al. 2019) were obtained from Dr. Per-Johan Jakobsson at Karolinska Institutet, Sweden. All the above compounds except L902688, which was dissolved in methanol, were dissolved in DMSO (Sigma, 472301) and stored at −80 °C in aliquots. Treatments of COCS with PGE2 (1 μM), PGD2 (1 μM), C118 (2 μM) and C934 (2 μM), 17-pt-PGE2 (1 μM and 5 μM) and L902688 (1 μM and 5 μM) were started from 14 days after the cultures were established and lasted to the end of experiments. COCS treated with same amounts of DMSO (or methanol for L902688) were used as controls. Recombinant human PDGFAA (110-13A) was purchased from Peprotech, dissolved in Opti-MEM, and stored at -20 °C in aliquots. Treatment of COCS with PDGFAA (40 ng/ml) was performed following the same protocol as with the compounds, except that COCS treated with Opti-MEM were used as controls.

### NG2 glia depletion

NG2 glia depletion in COCS and in vivo was performed as previously described (Liu and Aguzzi 2020). Briefly, to achieve ex vivo NG2 glia depletion, CP673451 (MedChemExpress, HY-12050) was supplemented in slice culture medium with the concentration of 1 μM. Treatments were started from 14 days after the cultures were established and lasted to the end of experiments (for long-term continuous depletion) or 10 days (for short-term transient depletion). Fresh CP673451 was added every time when the culture medium was changed. COCS treated with same amounts of DMSO were used as controls. To achieve in vivo NG2 glia depletion, tamoxifen-treated Pdgfrα^iDTR^ mice were injected intraperitoneally with diphtheria toxin (DT, Sigma, D0564) diluted in saline for five consecutive days (two injections per day with an 8-hour interval, 200 ng per injection). DT-injected tamoxifen-treated Pdgfrα-CreER mice and iDTR mice were pooled together and used as controls.

### Lentiviral production

Lenti-vectors used for expressing human PGE2 receptors EP1 (pLenti-PTGER1-mGFP-P2A-Puro, RC208597L4), EP2 (pLenti-PTGER2-mGFP-P2A-Puro, RC210883L4), EP3 (pLenti-PTGER3-mGFP-P2A-Puro, RC220173L4) and EP4 (pLenti-PTGER4-mGFP-P2A-Puro, RC210932L4) were purchased from OriGene Technologies. A control vector (pLenti-HygR-mGFP-P2A-Puro) was produced in house by replacing the EP2 sequence in the pLenti-PTGER2-mGFP-P2A-Puro plasmid with the sequence of the Hygromycin resistant gene (HygR). All plasmids were verified by Sanger sequencing before lentiviral production in HEK293T cells maintained in Opti-MEM supplemented with 10% fetal bovine serum (FBS). Briefly, HEK293T cells were seeded in 10-cm cell culture dishes and transfected at ∼ 80% confluency with a packaging plasmid mixture (transgene plasmid; VSVG plasmid; PAX2 plasmid) using FuGENE HD transfection reagent (Promega, E2311). Twenty-four hours after transfection, culture medium was changed to remove the transfection reagent. After another 48 hours, the culture medium containing lentivirus was collected, centrifuged at 1500g for 10 minutes and filtered through 0.45-micron Whatman filter units (GE Healthcare, 10462100). High-titer lentivirus was produced by concentrating the filtered supernatant with Lenti-X concentrator (Takara, 631231) and stored at -80 °C in aliquots.

### Primary microglia culture

Serum-free mouse microglia cultures were established using the immunopanning method according to previously published protocols with small modifications. Briefly, brain tissues (olfactory bulb, cerebellum and subcortical regions removed) from 5-day old C57BL/6J mice were dissected in cold Hanks’ Balanced Salt solution (HBSS, TermoFisher Scientific, 14175095) under a stereomicroscope. After removing meninges, brain tissues were minced and washed three times with HBSS. Afterwards, the minced brain tissues were digested in papain solution (Worthington Biochemical Corporation, LK003178) with DNase1 (Worthington Biochemical Corporation, LK003172) for 20 minutes at 37 °C and pipetted into single-cell suspension. The digestion solution was removed through centrifugation (1000g for 5 minutes), and cell pellets were resuspended in OptiMEM and filtered through a 70-μm cell strainer for immnopanning at room temperature. Each Petri dish used for immnopanning was coated with 30 μl goat antirat IgG (Jackson ImmunoResearch 112-005-167) diluted in 10 ml of sterile 50 mM Tris-HCl (pH 9.5) overnight at 4 °C. After 5 times of washing in phosphate buffered saline (PBS), the dishes were further coated with 30 μl rat anti-mouse CD11b antibody (ThermoFisher Scientific, 14-0112-82) diluted in the same buffer overnight at 4 °C and washed 5 times in PBS before use. After immunopanning for 45 minutes (gently swirl the dishes every 15 minutes), the floating cells in the suspension were removed from the dishes, and cells attached to the dishes were washed 5 times with PBS and digested with trypsin-EDTA (ThermoFisher Scientific, 25200056) for 10 minutes at 37 °C. Afterwards, the cells were collected by centrifugation, resuspended, and seeded in ibiTreat 8-well slide chambers (Ibidi, 80806) in OptiMEM-based medium containing 1% Sato Mix, 1% GlutaMax, 1% penicillin/streptomycin, 2 ng/ml human Tgfb2 (ThermoFisher Scientific, 100-35B), 10 ng/ml murine Csf1 (Biolegend, 576404) and 1.5 mg/ml cholesterol (Merck, C3045). After 3 days in culture, primary microglia were treated with recombinant human Tgfb2 (100 ng/ml), human Pleiotrophin (100 ng/ml, ThermoFisher Scientific, 450-15), human Midkine (100 ng/ml, ThermoFisher Scientific, 450-16), human Bmp7 (100 ng/ml, ThermoFisher Scientific, 120-03P) or human Semaphorin3d (400 ng/ml, Novus Biologicals, H00223117-P01) for 72 hours in the presence or absence of 0.01% w/v prion-containing brain homogenates, followed by PFA fixation and immunofluorescence.

### Primary neuronal culture

Primary neuronal cultures were established using cerebellar tissues from 5-day old C57BL/6J mice according to previously published protocols. Briefly, after removing meninges, cerebellar tissues were minced, washed three times with HBSS, digested in papain solution with DNase1 for 15 minutes at 37 °C and pipetted into single-cell suspension. After centrifugation, cell pellets were resuspended in OptiMEM and incubated in uncoated tissue culture dishes for 10 min to remove astrocytes and microglia. Afterwards, the cells were recollected, centrifuged, resuspended in neuronal culture medium containing Neurobasal medium (ThermoFisher Scientific, 21103049), 1% N2 (ThermoFisher Scientific, 17502048), 1% B27 (ThermoFisher Scientific, 17504044), 1% MEM Non-Essential Amino Acids (MEM NEAA, ThermoFisher Scientific, 11140050), 1% GlutaMax and 1% penicillin/streptomycin, and seeded in 48-well plates coated with PDL (ThermoFisher Scientific, A3890401). After 1 day, the cultures were treated with 2 μM AraC (British Pharmacopoeia, 383) for 24 hours to further remove glial cells. For prion infection, 0.01% w/v prion-containing brain homogenates were added into the cultures at day 5. After 10 days of prion infection, the cultures were treated with different concentrations of 17-pt-PGE2 and L902688 for 48 hours or controls (DMSO for 17-pt-PGE2 and methanol for L902688), followed by fixation and immunofluorescence.

### HovS cell culture

HovS cells (Avar, Heinzer et al. 2020), a subclone of the human SH-SY5Y cell line, where the human PRNP gene was replaced with the ovine PRNP VRQ allele, were maintained in OptMEM medium supplemented with 10% FBS, 1% MEM-NEAA, 1% GlutaMax, 1% penicillin/streptomycin and 400 μg/ml Geneticin (ThermoFisher Scientifc, 10131027). To establish a cellular model of chronic prion infection, HovS cells were exposed to brain homogenates derived from PG127-prion-infected tg338 mice and passaged for at least 10 times before being used for experiments. To investigate the effects of PGE2 and the involved PGE2 receptors in prion-induced cell toxicity, PG127-HovS cells were seeded into 96-well plates. Twenty-four hours after seeding, cells were transduced with either control lentivirus or lentivirus harboring one of the four EP receptor transgenes. After 48 hours, the lentivirus-containing culture medium was removed, and fresh culture medium supplemented with DMSO or PGE2 (10 mM) was added in the wells and incubated for another 48 hours. Finally, GFP^+^ cells (cells infected by lentivirus) were imaged with a Nikon Eclipse Ti2-E fluorescent microscope and quantified with ImageJ.

### Western blotting

Western blotting was performed as previously described (Liu, Sorce et al. 2018). Briefly, cultured brain slices or collected brain tissues were lysed with a bead-mill homogenizer in RIPA buffer supplemented with proteinase inhibitor cocktail cOmplete (MERCK, 11697498001) and phosphatase inhibitor cocktail PhosSTOP (MERCK, 4906845001). After centrifuge, protein concentrations in the supernatant were quantified with the BCA method. Protein samples were then mixed with western blotting loading buffer and boiled for 5 minutes at 95 °C before being loaded onto gels. For western blotting aimed to detect prions, samples were first digested with 10 μg/ml (for COCS) or 20 μg/ml (for brain tissues) proteinase K for 30 minutes at 37 °C, then mixed with western blotting loading buffer and boiled for 5 minutes at 95 °C. The following primary antibodies were used: mouse monoclonal antibody against actin (1:10,000, Merck Millipore, MAB1501R); mouse monoclonal antibody against PrP (POM1, 1:5000, homemade); rabbit polyclonal antibody against NG2 (1:500, MERCK, AB5320); rabbit polyclonal antibody against PDGFRα (1:500, Santa Cruz, sc-338); rabbit monoclonal antibody against NeuN (1:2000, Abcam, ab177487). Depending on the primary antibodies used, suitable HRP-conjugated secondary antibodies (1:10,000, Jackson ImmunoResearch Laboratories) were chosen. Membranes were developed with Crescendo Western HRP Substrate (MERCK, WBLUR0500), visualized, and digitized with ImageQuant (LAS-4000; Fujifilm, Japan). Optical densities of bands were analyzed using ImageJ.

### Immunofluorescence

Immunofluorescent staining of cultured brain slices, primary cells and cryosections was performed according to procedures published previously (Liu and Aguzzi 2020). For cultured brain slices: after removing culture medium, brain slices were washed with PBS and fixed in 4% PFA for 30 min at room temperature. After several washes to remove residual PFA, brain slices were then permeabilized with PBST (PBS + 0.1% Triton X-100) for 2 hours at room temperature and blocked with 5% goat serum (GS) overnight at 4 °C before adding primary antibodies. For primary cells: after removing culture medium, cells were washed with PBS and fixed in 4% PFA for 30 min at room temperature. After several washes to remove residual PFA, cells were blocked with 5% GS for 2 hours at room temperature before adding primary antibodies. For cryosections: mice were perfused transcardially with 20 ml PBS and 20 ml 4% PFA. The dissected brains were then postfixed with 4% PFA for 4 hr or overnight in the fridge. After removing PFA and brief washing with PBS, brains were transferred into 30% sucrose solution for dehydration, and kept at 4°C. After dehydration was complete, brains were cut into 25-μm thick cryosections. To perform immunostaining, cryosections were permeabilized in PBST for 2 hours and blocked with 5% GS for 2 hours at room temperature before incubation in primary antibodies. The following primary antibodies were used: rabbit polyclonal antibody against NG2 (1:500, a gift from Prof. Stallcup), rabbit monoclonal antibody against NeuN (1:1000, Abcam, ab177487), rabbit polyclonal antibody against Iba1 (1:500, Wako, 019-19741), rat monoclonal antibody against Cd68 (1:200, BioRad, MCA1957), rabbit polyclonal antibody against Map2 (1:200, Biolegend, 840601), mouse monoclonal antibody against Tau (1:200, ThermoFisher Scientific, MN1010), chicken polyclonal antibody against NeuN (1:1000, Merck, ABN91), mouse monoclonal antibody against Cox2 (1:200, Santa Cruz, sc-166475), mouse monoclonal antibody against Ptges (1:200, Santa Cruz, sc-365844), rabbit polyclonal antibody against EP1 (1:200, Bioss Antibodies, BS-6316R), rabbit monoclonal antibody against EP2 (1:200, Abcam, ab167171), rabbit polyclonal antibody against EP3 (1:200, Cayman Chemical, 101760) and mouse monoclonal antibody against EP4 (1:200, ProteinTech, 66921-1-Ig). Cultured brain slices were incubated in primary antibody at 4°C for 3 days. Cryosections and primary cells were incubated in primary antibody at 4°C overnight. After several washes in PBST, cultured brain slices were incubated in suitable secondary antibodies overnight at 4°C, and cryosections and primary cells were incubated in suitable secondary antibodies for 2 hours at room temperature. Immunofluorescent images were captured using a FLUOVIEW FV10i confocal microscope (Olympus Life Science) or Nikon Eclipse Ti2-E fluorescent microscope, and were quantified using ImageJ.

### Quantitative real-time PCR

To perform quantitative real-time-PCR (qRT-PCR) analysis, total RNA from cultured brain slices or collected brain tissues was extracted using TRIzol reagent (ThermoFisher Scientific, 15596026). QuantiTect Reverse Transcription kit (QIAGEN, 205311) was used to synthesize cDNA from the extracted RNA samples. qRT-PCR was performed on a ViiA7 Real-Time PCR system (Applied Biosystems, USA) using the SYBR Green PCR Master Mix (ThermalFisher Scientifc, 4309155). We used the following primers for qRT-PCR analysis: Mouse actin: sense, 5′-AGATCAAGATCATTGCTCCTCCT-3′, antisense, 5′-ACGCAGCTCAGTAACAG-TCC-3′. Mouse NG2: sense, 5′-ACCCAGGCTGAGGTAAATGC-3′, antisense, 5′- ACAGGCAGCATCGAAAGACA-3′. Mouse Pdgfrα: sense, 5′-ATTAAGCCGGTCCCAACCTG-3′, antisense, 5′-AATGGGACCTGACTTGGTGC-3′. Mouse TNFα: sense, 5′-ACGTCGTAG- CAAACCACCAA-3′, antisense, 5′-ATAGCAAATCGGCTGACGGT-3′. Mouse IL1β: sense, 5′- TGCAGCTGGAGAGTGTGGATCCC-3′, antisense, 5′-TGTGCTCTGCTTGTGAGGTGCTG-3′. Mouse IL12β: sense, 5′-TGGTTTGCCATCGTTTTGCTG-3′, antisense, 5′- ACAGGTGAGGTTCACTGTTTCT-3′. Mouse Cox1: sense, 5′-TCCATCCACTCCCAGAG-TCAT-3′, antisense, 5′-ACAACAGGGATTGACTGGTGA-3′. Mouse Cox2: sense, 5′- GGGCCATGGAGTGGACTTAAA-3′, antisense, 5′-ACTCTGTTGTGCTCCCGAAG-3′. Mouse Ptges: sense, 5′-TCTCACTCTCAGTCCCGGTG-3′, antisense, 5′-GGGGTTGGCAAAA- GCCTTC-3′. Mouse Ptgds: sense, 5′-GCTCCTTCTGCCCAGTTTTCC-3′, antisense, 5′- CCCCAGGAACTTGTCTTGTTGA-3′.

### Single-cell RNA sequencing analysis

Cerebral cortex and hippocampal single-cell RNA sequencing dataset from prion-infected or control mice (Slota, Sajesh et al. 2022) were obtained from the single cell portal at the Broad institute through the following link: https://singlecell.broadinstitute.org/single_cell/study/SCP1962. After loading the raw single-cell RNA sequencing data into R, doublets and ambient RNA were removed using scrublet (Wolock, Lopez et al. 2019) and decontX (Yang, Corbett et al. 2020), respectively. Cells with nFeature_RNA less than 1000 or more than 7000 or mitochondrial RNA percentage more than 5% were filtered out using Seurat (Stuart, Butler et al. 2019). After normalizing the data and regressing out the mitochondrial genes and cell cycle effects with Seurat, data from different animals were integrated using Harmony (Korsunsky, Millard et al. 2019). After data integration, cells were clustered using UMAP, and the resulting clusters were annotated based on known cell-type markers. To molecularly characterize Cox2^+^ and Ptges^+^ microglia populations in the prion-infected mouse brain, microglia clusters were selected for further analysis, and differentially expressed genes between the Cox2^+^ and Cox2^-^ or Ptges^+^ and Ptges^-^ microglia were identified using the FindMarkers function in Seurat based on the cutoff FDR < 0.05 and logfc.threshold > 0.25. Changes of cell-cell interactions between NG2 glia and Cox2^+^ or Cox2^-^ microglia at the single-cell level in the prion-infected mouse brain were analyzed using the CellChat package (Jin, Guerrero-Juarez et al. 2021).

### Statistical analyses

Unless otherwise mentioned, unpaired, two-tailed student’s t test was used for comparing data from two groups. All data were presented as mean ± SEM. To compare the incubation time of prion-inoculated mice, survival curves were estimated using the Kaplan-Meier method and compared statistically using the log rank test. Statistical analysis and data visualization were done using GraphPad Prism 9. P-value < 0.05 was considered statistically significant.

## Acknowledgements

We thank Dr. William Stallcup for providing the polyclonal antibodies against NG2, Julie Domange, Mirzet Delic, Maria Svercelova and Marija Mihailova for excellent assistance. YL was supported by a Forschungskredit PostDoc grant (FK Liu 19-042). AA was supported by an Advanced Grant of the European Research Council (ERC 670958), the Swiss National Foundation (SNF 179040), and the Nomis Foundation.

## Author Contributions

YL and AA conceptualized the study. YL designed and performed the experiments with help from JG, MM and MK. PJ provided the Ptges inhibitors and helpful guidance on relevant experiments. YL and AA wrote the manuscript.

## Liu et al., Supplementary figures

**Supplementary figure 1.**
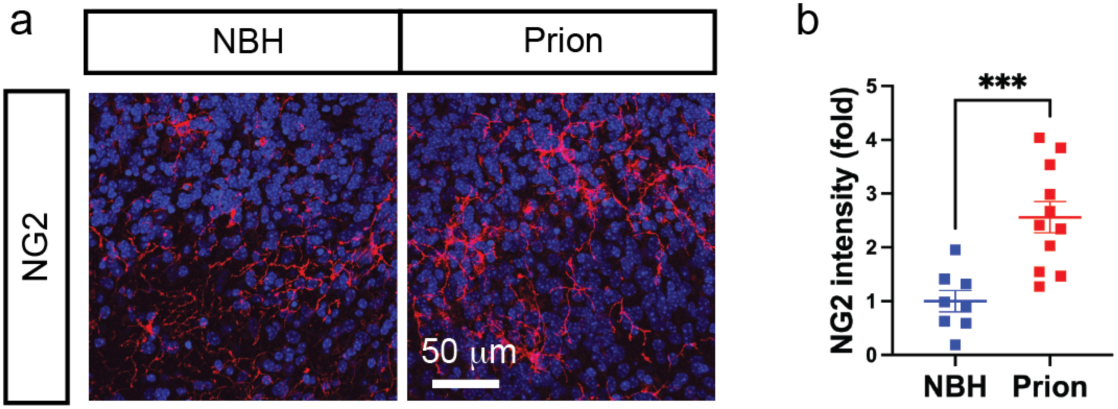
**a**, NG2 immunofluorescence showing NG2 glia activation in prion-infected C57BL/6J COCS vs C57BL/6J COCS exposed to NBH. Nuclei were stained with DAPI (blue). **b**, Quantification of NG2 immunointensity shown in **a**. n = 8 slices for NBH; n = 11 slices for Prion. Here and in following figures: *: p < 0.05; **: p < 0.01; ***: p < 0.001; n.s: not significant.

**Supplementary figure 2.**
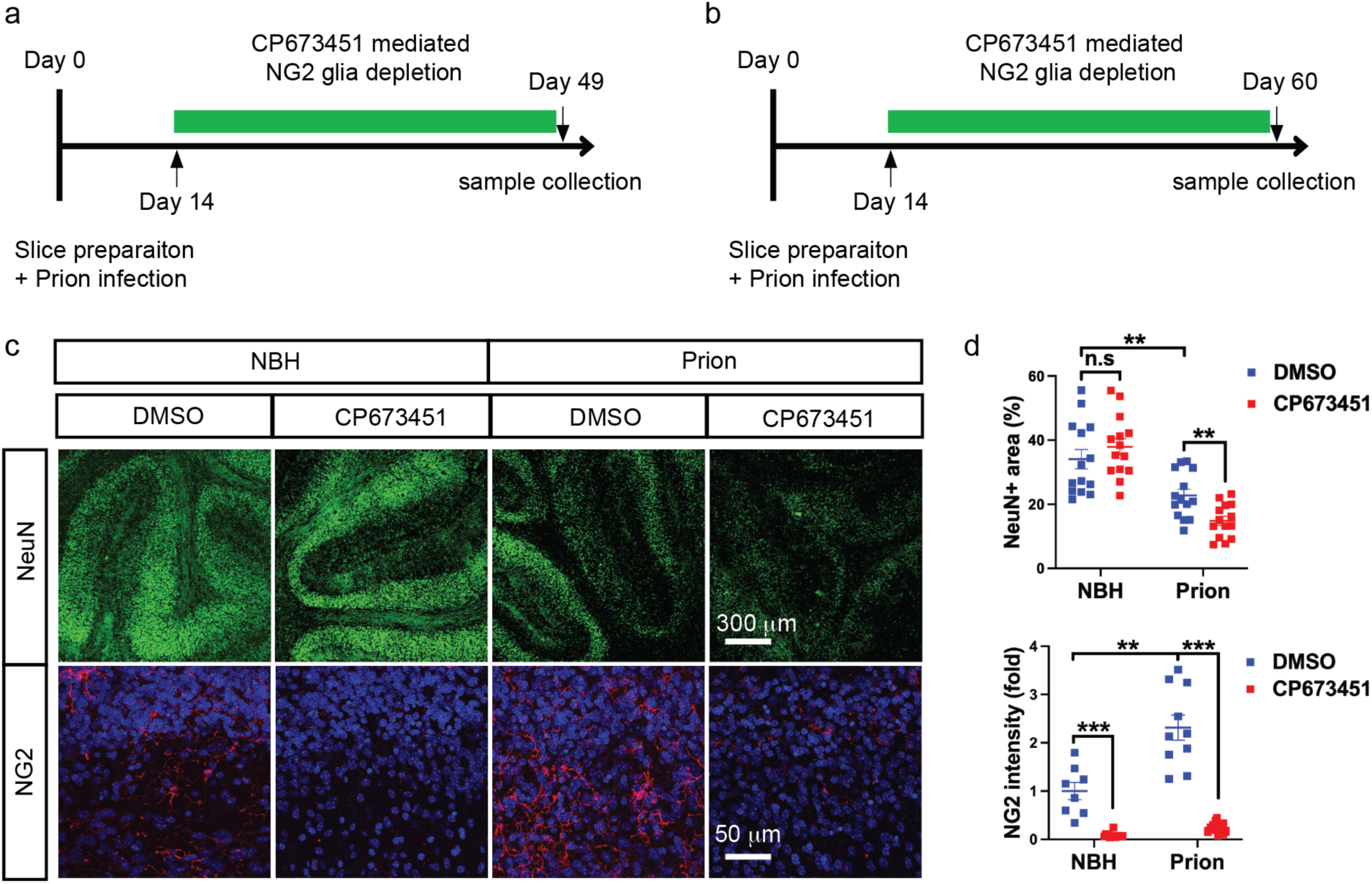
**a-b**, Experimental design of NG2 glia depletion in Tga20 COCS (**a**) and C57BL/6J COCS (**b**). **c**, NeuN and NG2 immunofluorescence showing enhanced neurodegeneration in continuously NG2-glia-depleted (CP673451) C57BL/6J COCS compared to NG2 glia intact (DMSO) C57BL/6J COCS after prion infection. Nuclei were stained with DAPI (blue). **d**, Quantification of NG2 immunointensity and NeuN positive area shown in **c**. n = 14 slices/condition for NeuN and n > 8 slices/condition for NG2.

**Supplementary figure 3.**
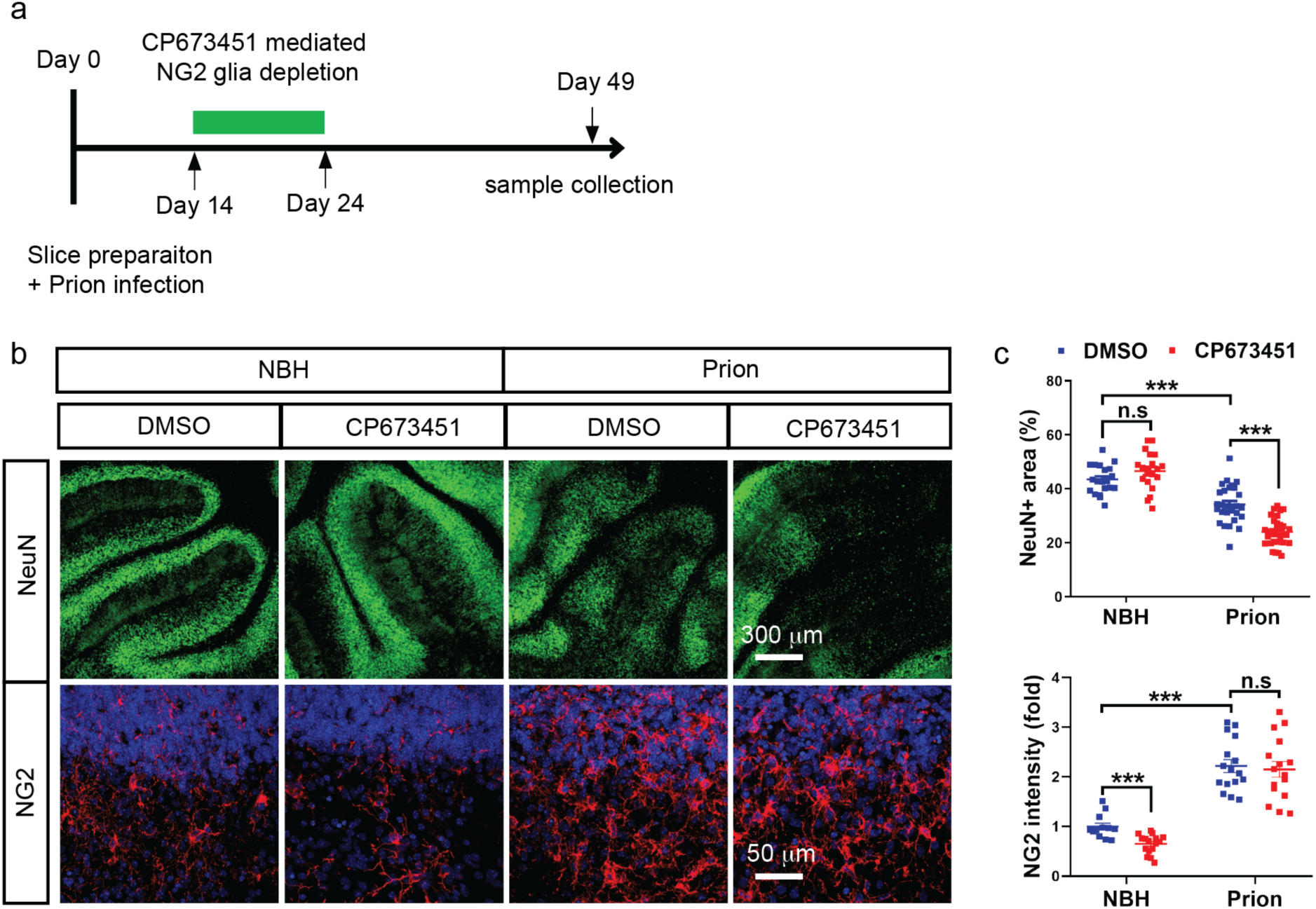
**a**, Experimental design of transient NG2 glia depletion in Tga20 COCS. **b**, NeuN and NG2 immunofluorescence showing enhanced neurodegeneration in transiently NG2-glia-depleted (CP673451) Tga20 COCS compared to NG2 glia intact (DMSO) Tga20 COCS after prion infection. NG2 glia density is largely recovered in prion-infected, transiently NG2-glia-depleted COCS at the end of the experiment. Nuclei were stained with DAPI (blue). **c**, Quantification of NG2 immunointensity and NeuN positive area shown in **b**. n > 16 slices/condition.

**Supplementary figure 4.**
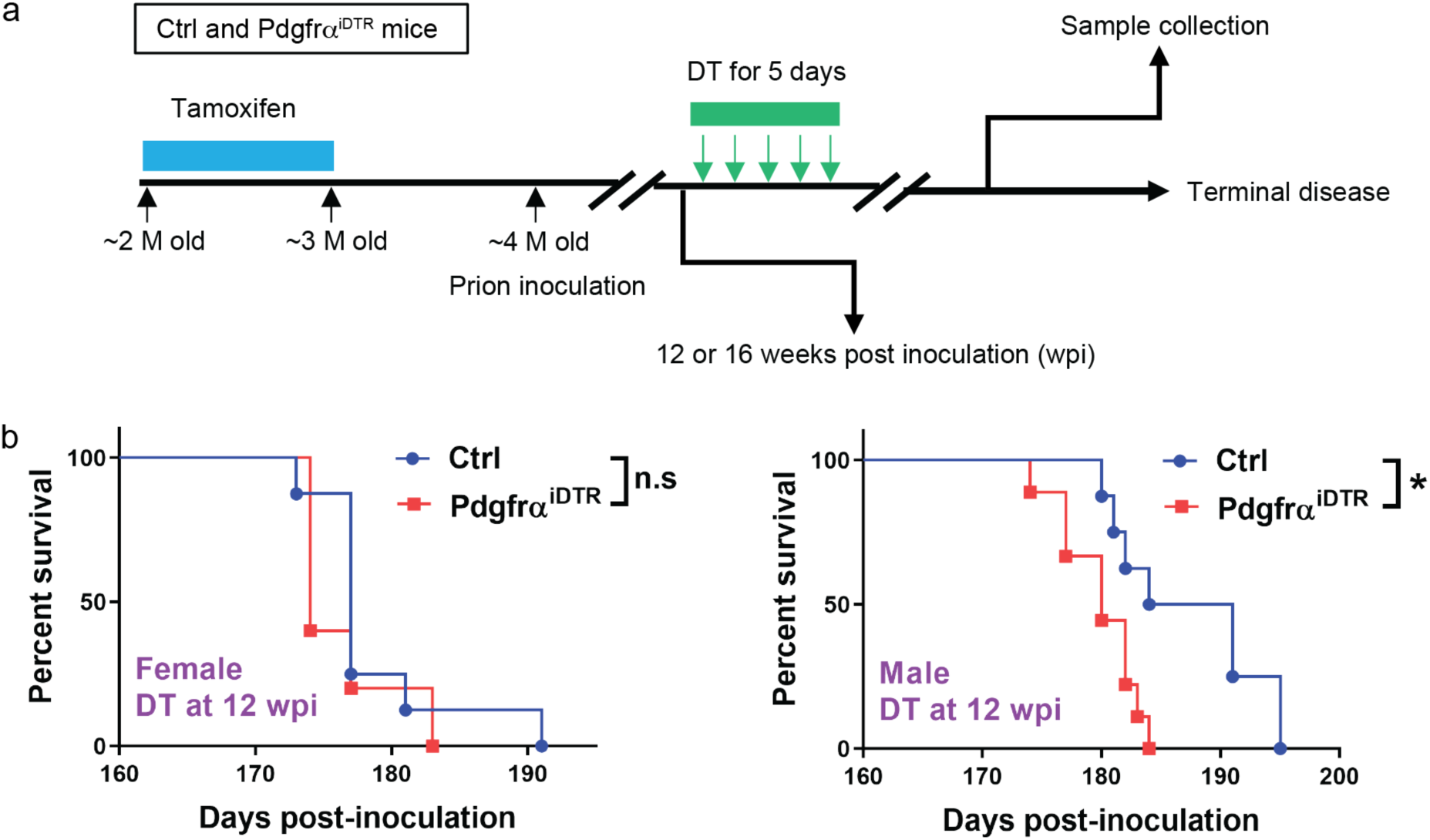
**a**, Experimental design of NG2 glia depletion in prion inoculated Pdgfrα^iDTR^ mice. Littermate Pdgfrα-CreER and iDTR mice were pooled together as control (Ctrl) and treated with tamoxifen and DT the same way as the Pdgfrα^iDTR^ mice. **b**, Survival curves showing accelerated prion disease in male, but not female NG2-glia-depleted (Pdgfrα^iDTR^) mice compared to NG2 glia intact (Ctrl) mice after prion inoculation (median survival: 180 days for male Pdgfrα^iDTR^ mice; 187.5 days for male control mice; 174 days for female Pdgfrα^iDTR^ mice; 177 days for female control mice). NG2 glia depletion was induced at 12 wpi.

**Supplementary figure 5.**
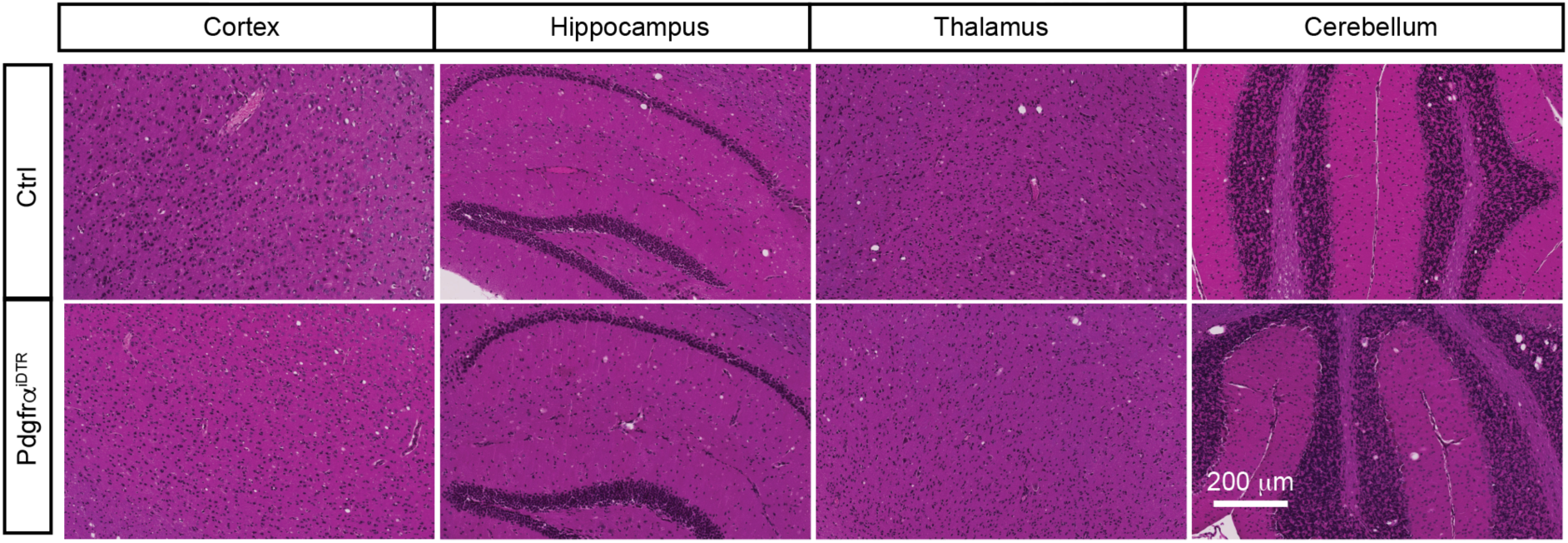
HE staining of brain tissues of NG2-glia-depleted (Pdgfrα^iDTR^) and NG2 glia intact (Ctrl) mice after prion inoculation indicating similar vacuolation in different brain regions. NG2 glia depletion was induced at 16 wpi.

**Supplementary figure 6.**
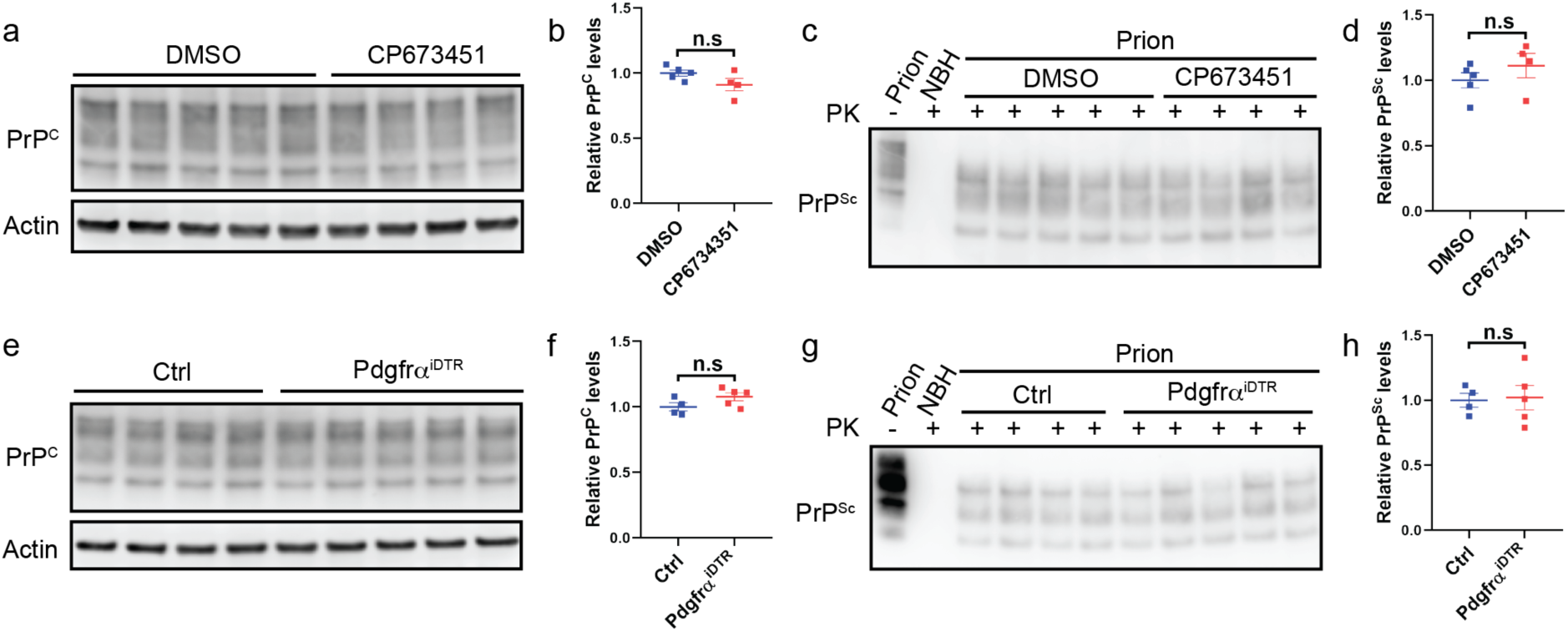
**a-b**, Western blots (**a**) and quantification (**b**) showing no changes of PrP^C^ level in NG2-glia-depleted (CP673451) Tga20 COCS compared to NG2 glia intact (DMSO) Tga20 COCS. n = 5 samples for DMSO and n = 4 samples for CP673451; 6 to 8 slices/sample. **c-d**, Western blots (**c**) and quantification (**d**) showing no changes of PK-resistant PrP^Sc^ levels in NG2-glia-depleted (CP673451) Tga20 COCS compared to NG2 glia intact (DMSO) Tga20 COCS after prion infection. n = 5 samples for DMSO and n = 4 samples for CP673451; 6 to 8 slices/sample. **e-f**, Western blots (**e**) and quantification (**f**) showing no changes of PrP^C^ levels in the brains of NG2-glia-depleted (Pdgfrα^iDTR^) mice compared to NG2 glia intact (Ctrl) mice. n = 4 mice for Ctrl and n = 5 mice for Pdgfrα^iDTR^. **g-h**, Western blots (**g**) and quantification (**h**) showing no changes of PK-resistant PrP^Sc^ levels in the brains of NG2-glia-depleted (Pdgfrα^iDTR^) mice compared to NG2 glia intact (Ctrl) mice after prion infection. NG2 glia depletion was induced at 16 wpi. n = 4 mice for Ctrl and n = 5 mice for Pdgfrα^iDTR^.

**Supplementary figure 7.**
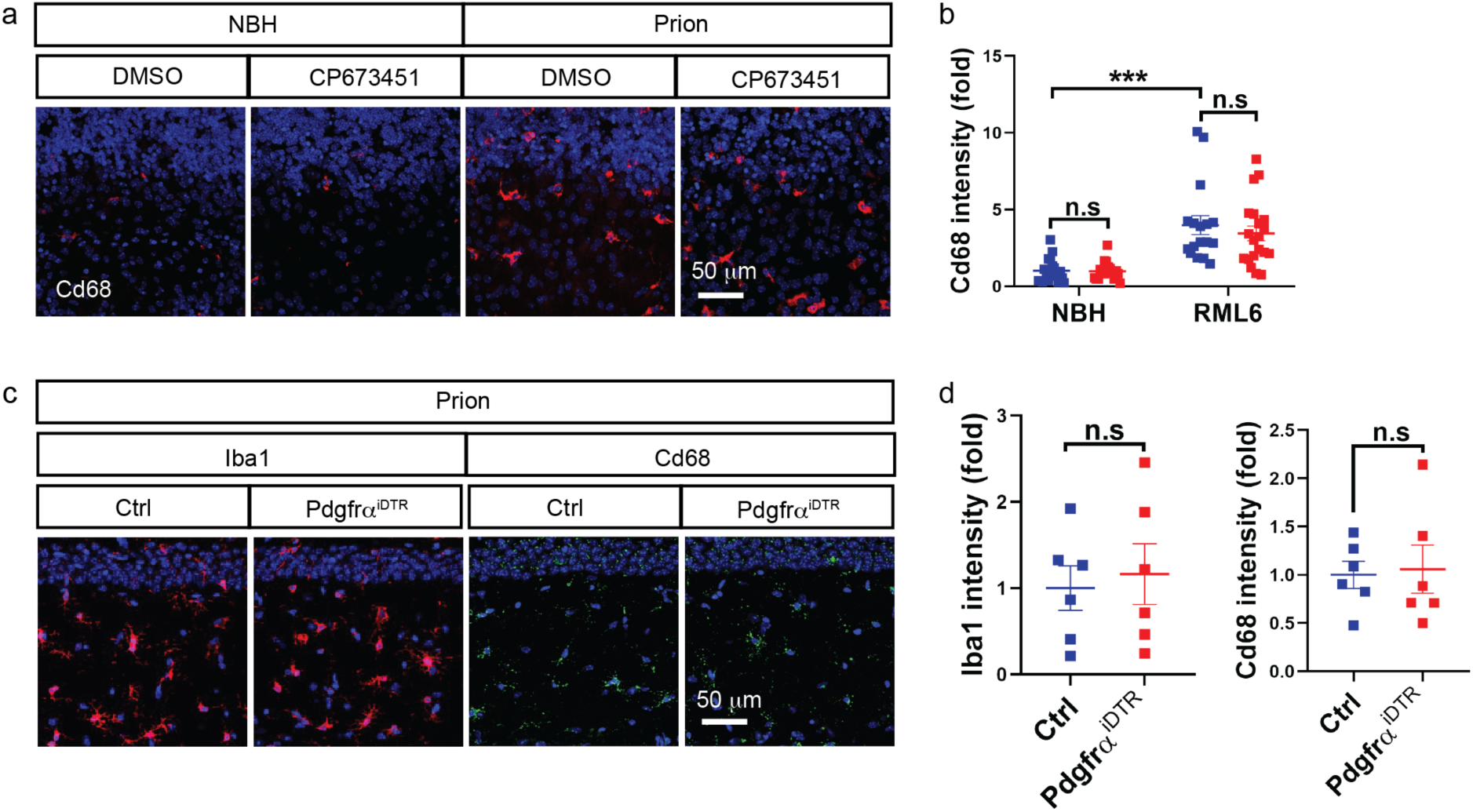
**a-b**, Cd68 immunofluorescence (**a**) and quantification (**b**) showing similar levels of microglia activation in NG2-glia-depleted (CP673451) Tga20 COCS compared to NG2 glia intact (DMSO) Tga20 COCS after prion infection. Nuclei were stained with DAPI (blue). n > 16 slices/condition. **c-d**, Iba1 and Cd68 immunofluorescence (**c**) and quantification (**d**) showing similar levels of microglia activation in the hippocampi of NG2-glia-depleted (Pdgfrα^iDTR^) mice compared to NG2 glia intact (Ctrl) mice after prion infection. NG2 glia depletion was induced at 16 wpi. Nuclei were stained with DAPI (blue). n = 6 mice/group.

**Supplementary figure 8.**
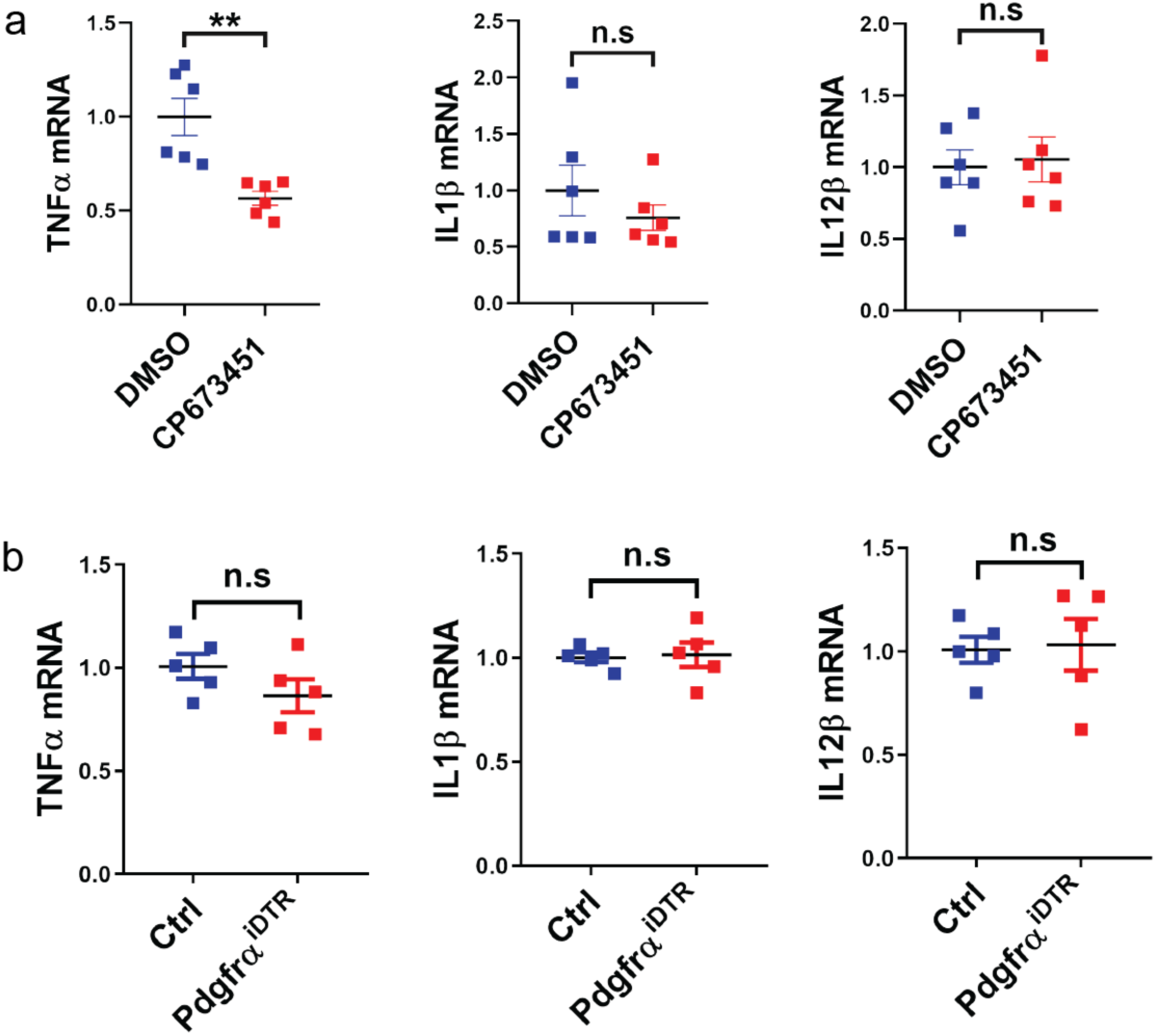
**a**, qRT-PCR results showing minimal changes of proinflammatory factors TNFα, IL1β and IL12β in NG2-glia-depleted (CP673451) Tga20 COCS compared to NG2 glia intact (DMSO) Tga20 COCS after prion infection. n = 6 samples/condition; 6 to 8 slices/sample. **b**, qRT-PCR results showing no changes of proinflammatory factors TNFα, IL1β and IL12β in the brains of NG2-glia-depleted (Pdgfrα^iDTR^) mice compared to NG2 glia intact (Ctrl) mice after prion inoculation. n = 5 mice/group. NG2 glia depletion was induced at 16 wpi.

**Supplementary figure 9.**
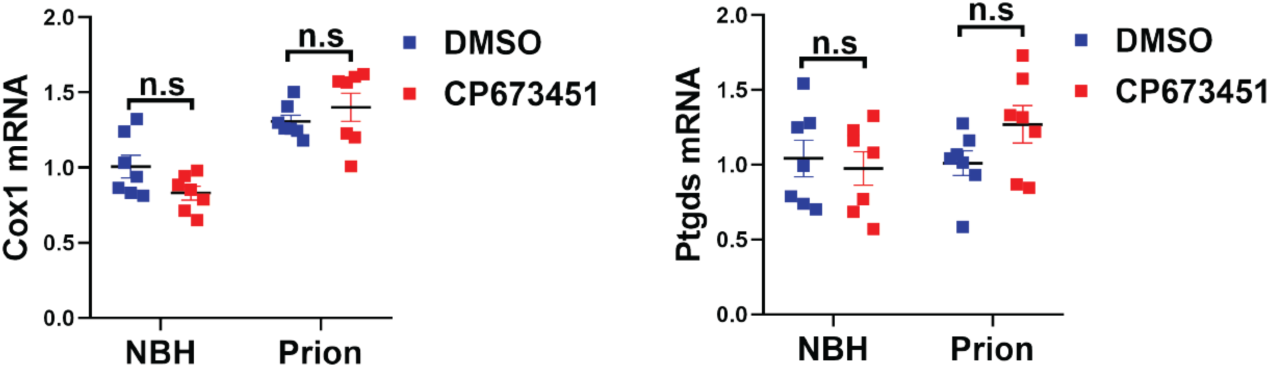
qRT-PCR results showing no changes of Cox1 and Ptgds in NG2-glia-depleted (CP673451) Tga20 COCS compared to NG2 glia intact (DMSO) Tga20 COCS in the presence or absence of prions. n = 7 samples/condition; 6 to 8 slices/sample.

**Supplementary figure 10.**
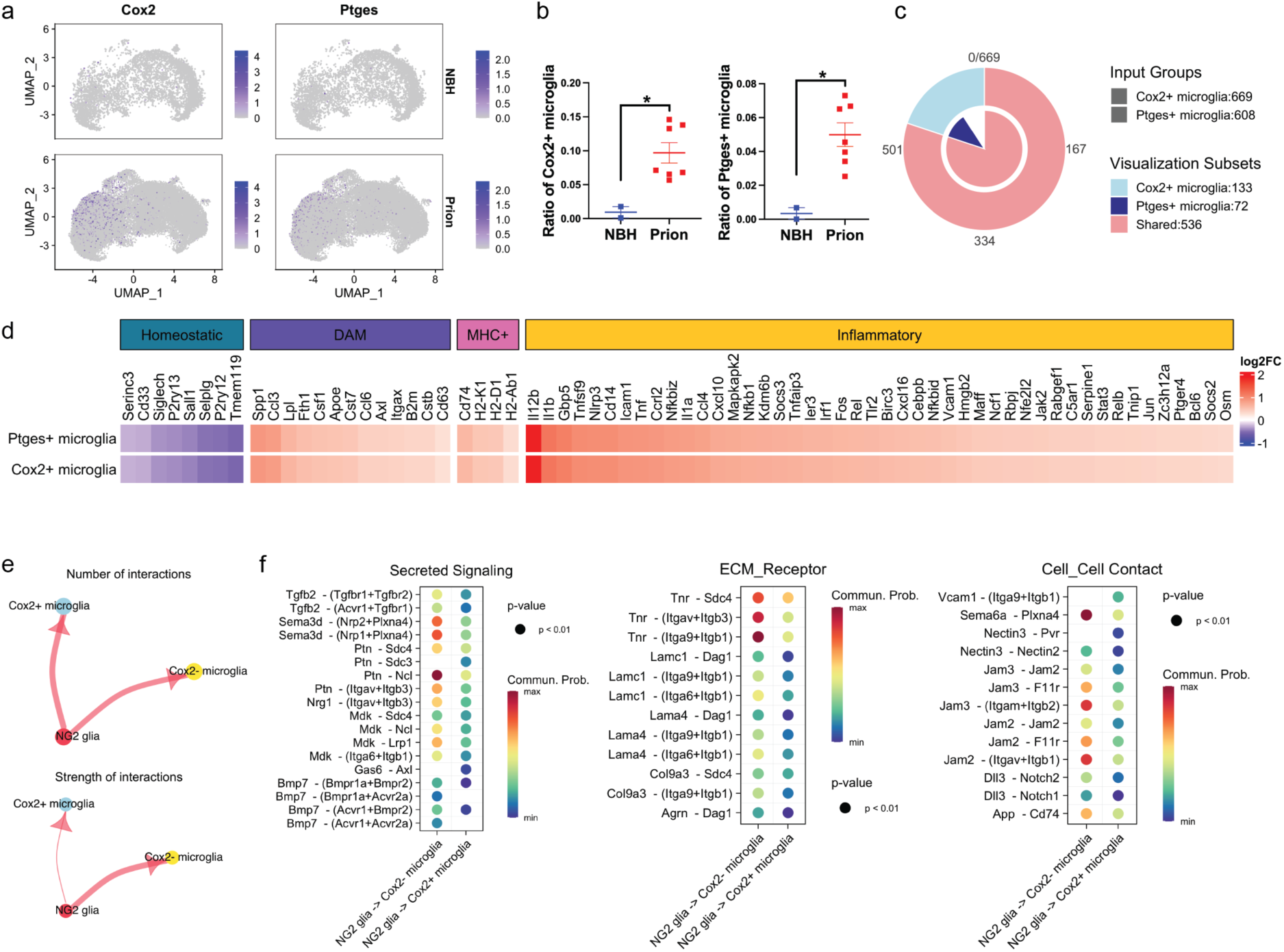
**a**, UMAP of single-cell RNA-seq data showing Cox2^+^ and Ptges^+^ microglia among total microglia in the hippocampus of prion- or NBH-inoculated mice. **b**, Quantification of Cox2^+^ and Ptges^+^ microglia fractions against total microglia in the hippocampus of prion- or NBH-inoculated mice shown in **a**. n = 2 mice for NBH and n = 7 mice for Prion. **c**, Venn diagram showing numbers of shared and distinct DEGs of Cox2^+^ and Ptges^+^ microglia in the hippocampus of prion-inoculated mice. **d**, Heatmap showing downregulation of homeostatic signature genes and upregulation of DAM and MHC^+^ microglia signature genes as well as inflammatory genes in Cox2^+^ and Ptges^+^ microglia in the hippocampus of prion-inoculated mice. **e**, CellChat analysis of cell-cell communications showing unaltered number of interactions (plotted as the thickness of the edges) but reduced strength of interactions (plotted as the thickness of the edges) from NG2 glia to Cox2^+^ microglia in the hippocampus of prion-inoculated mice. **f**, Heatmaps showing significantly weakened NG2 glia to Cox2^+^ microglia interaction pathways in the hippocampus of prion-inoculated mice.

**Supplementary figure 11.**
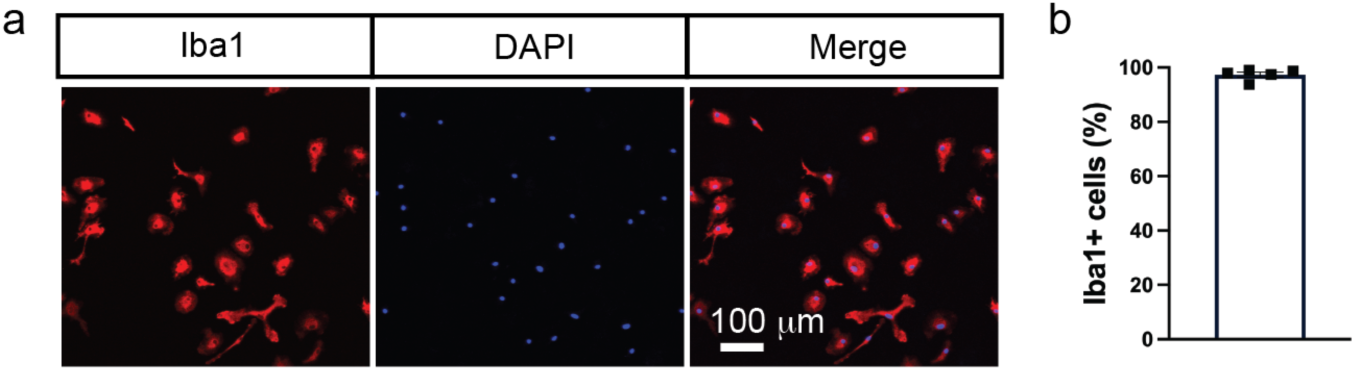
**a**, Iba1 immunofluorescence showing high purity of primary microglia cultures isolated by Cd11b immunopanning. Nuclei were stained with DAPI (blue). **b**, Quantification of Iba1+ cell percentage shown in **a**.

**Supplementary figure 12.**
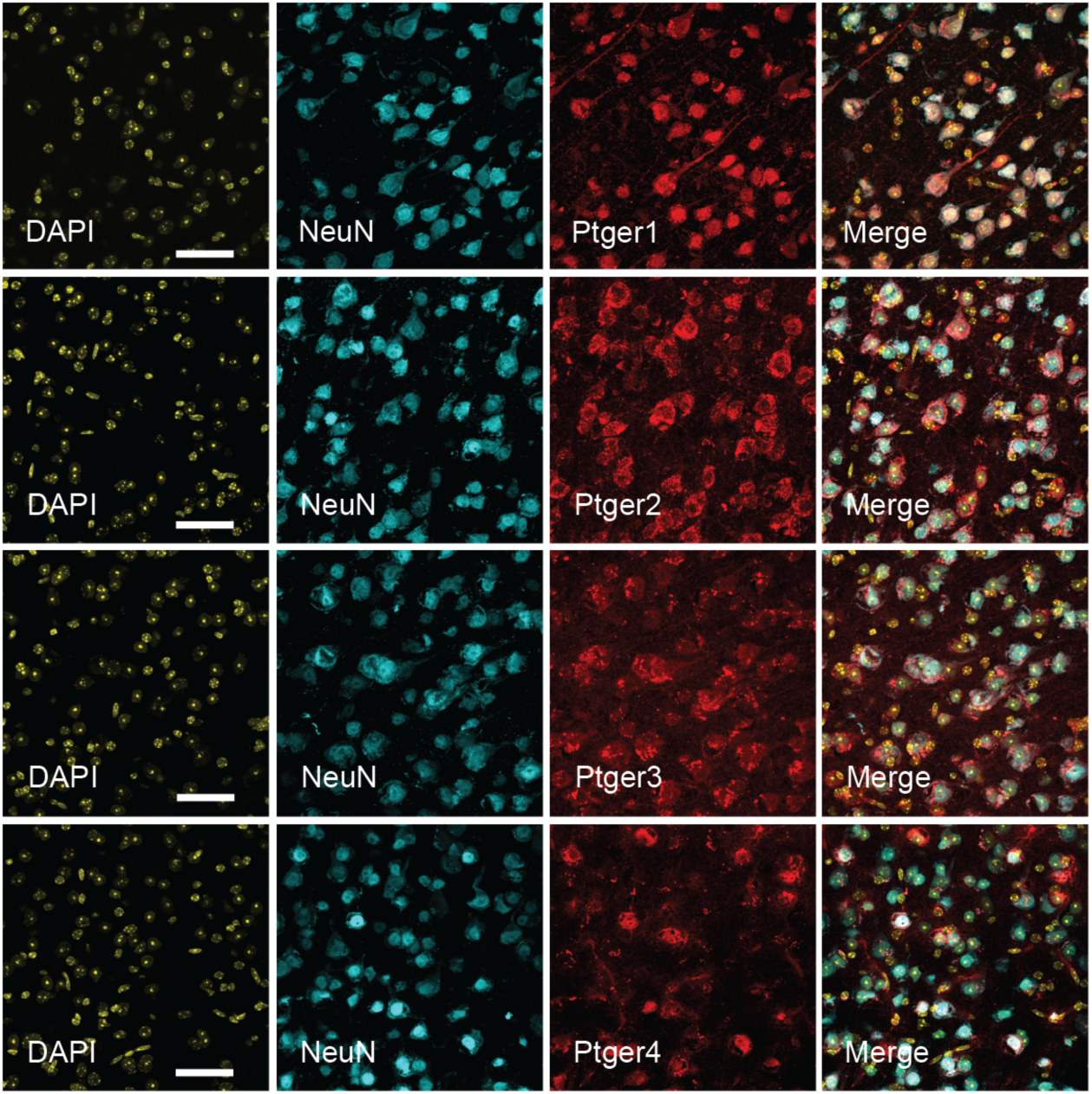
Immunofluorescence of NeuN and PGE2 receptors in the cerebral cortex showing neuronal expression of Ptger1, Ptger2, Ptger3 and Ptger4 in the adult mouse brain. Scale bar: 50 μm.

**Supplementary figure 13.**
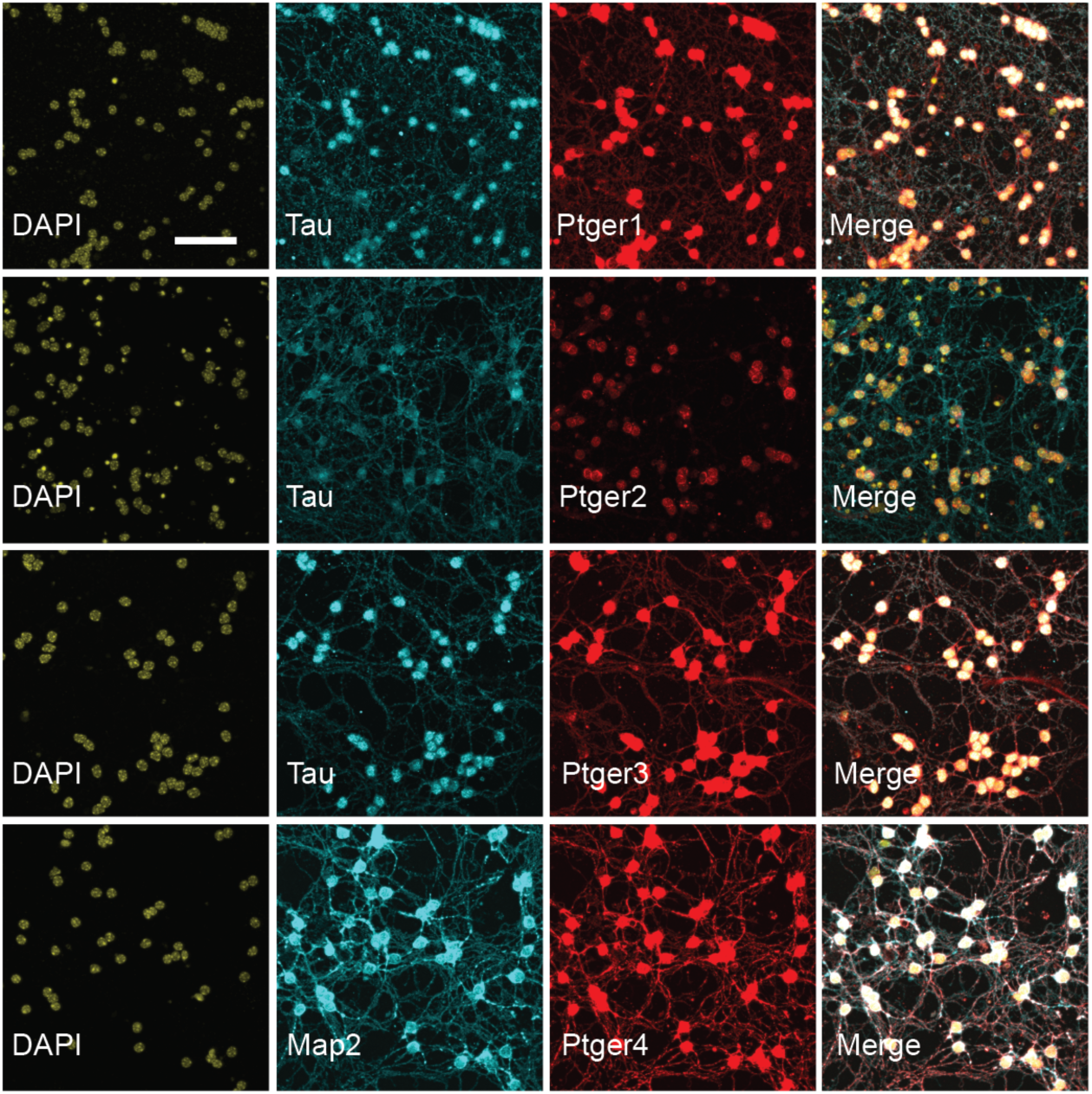
Immunofluorescence showing expression of Ptger1, Ptger2, Ptger3 and Ptger4 in the primary neuronal cultures (3 weeks in vitro). Ptger2 is mainly expressed in the neuronal cell body; Ptger1, Ptger3 and Ptger4 are expressed in both the neuronal cell body and neuronal processes. Scale bar: 50 μm.

**Supplementary figure 14.**
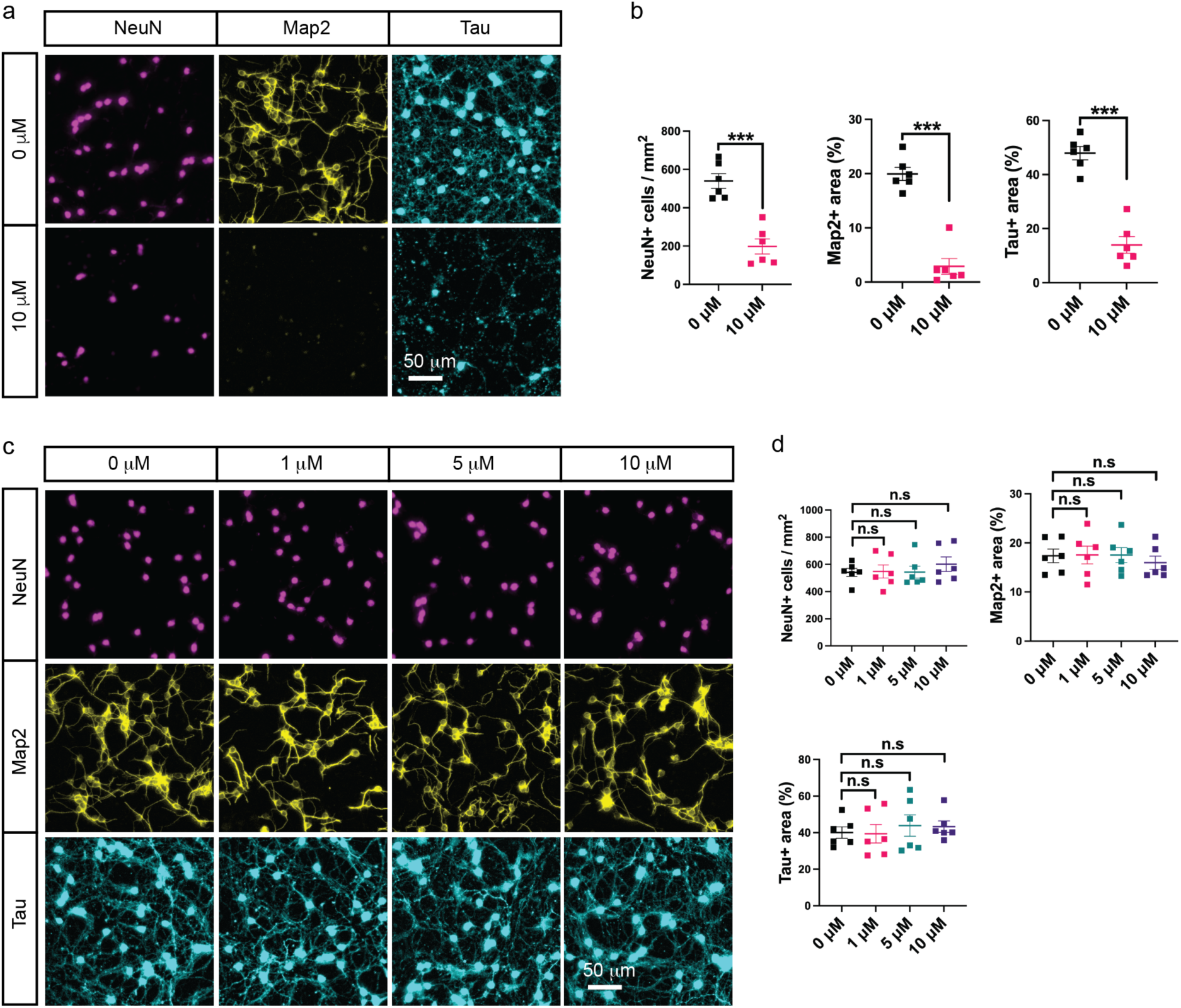
**a**, Immunofluorescence of NeuN, Map2 and Tau showing cellular damages of primary neurons treated with high concentration of Ptger4 agonist L902688. **b**, Quantification of neuronal density as well as Map2 positive and Tau positive areas shown in **a**. n = 6 independent experiments. **c**, Immunofluorescence of NeuN, Map2 and Tau showing no damages of prion-infected primary neurons treated with different concentrations of Ptger1 agonist 17-pt-PGE2. **d**, Quantification of neuronal density as well as Map2 positive and Tau positive areas shown in **c**. n = 6 independent experiments.

**Supplementary figure 15.**
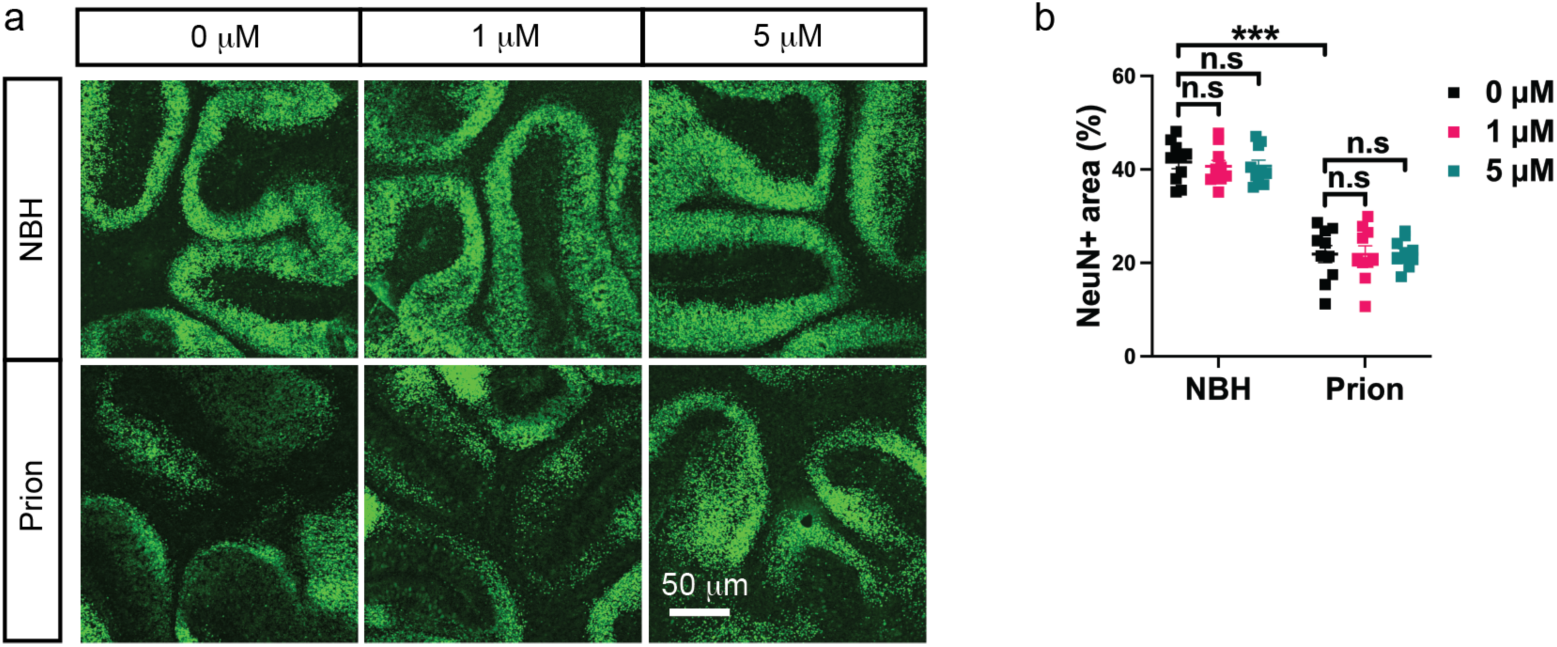
**a-b**, NeuN immunofluorescence (**a**) and quantification (**b**) showing no changes of prion-induced neurodegeneration in 17-pt-PGE2-treated COCS. n = 10 slices/condition.

**Supplementary figure 16.**
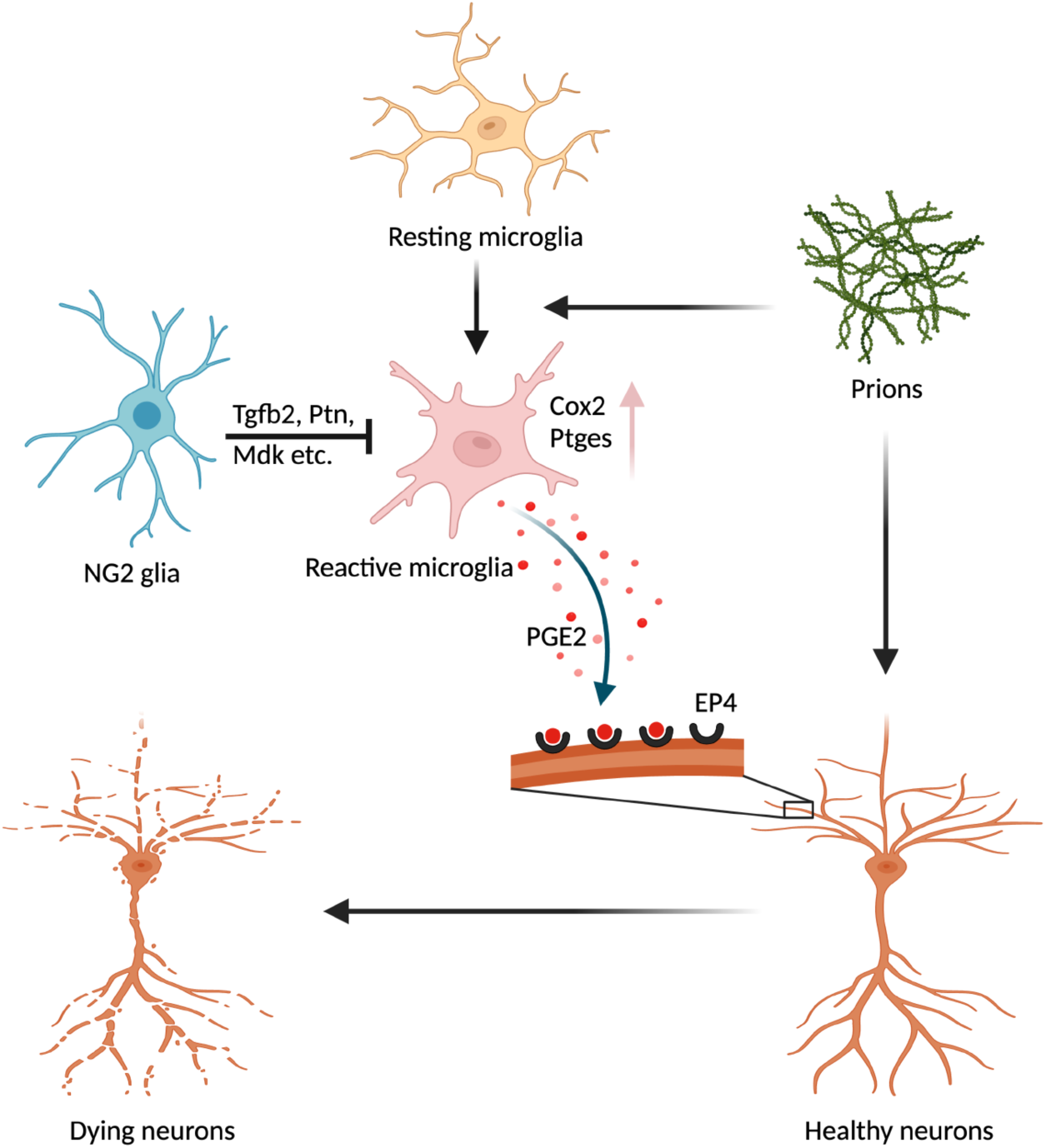
Diagram summarizing the main findings. In prion diseases, microglia become activated, and upregulate the pathway responsible for PGE2 biosynthesis, which promotes prion-induced neurodegeneration through binding to neuronal EP4 receptor. NG2 glia serve as a brake in this process, inhibiting microglial Cox2-Ptges pathway and PGE2 biosynthesis through multiple mechanisms (e.g., secreted signaling, ECM-receptor interaction and cell-cell contact). Several NG2-glia-derived factors playing a role in this process, such as Tgfb2, Pleiotrophin (Ptn) and Midkine (Mdk) are highlighted in the diagram.

**Supplementary figure 17.**
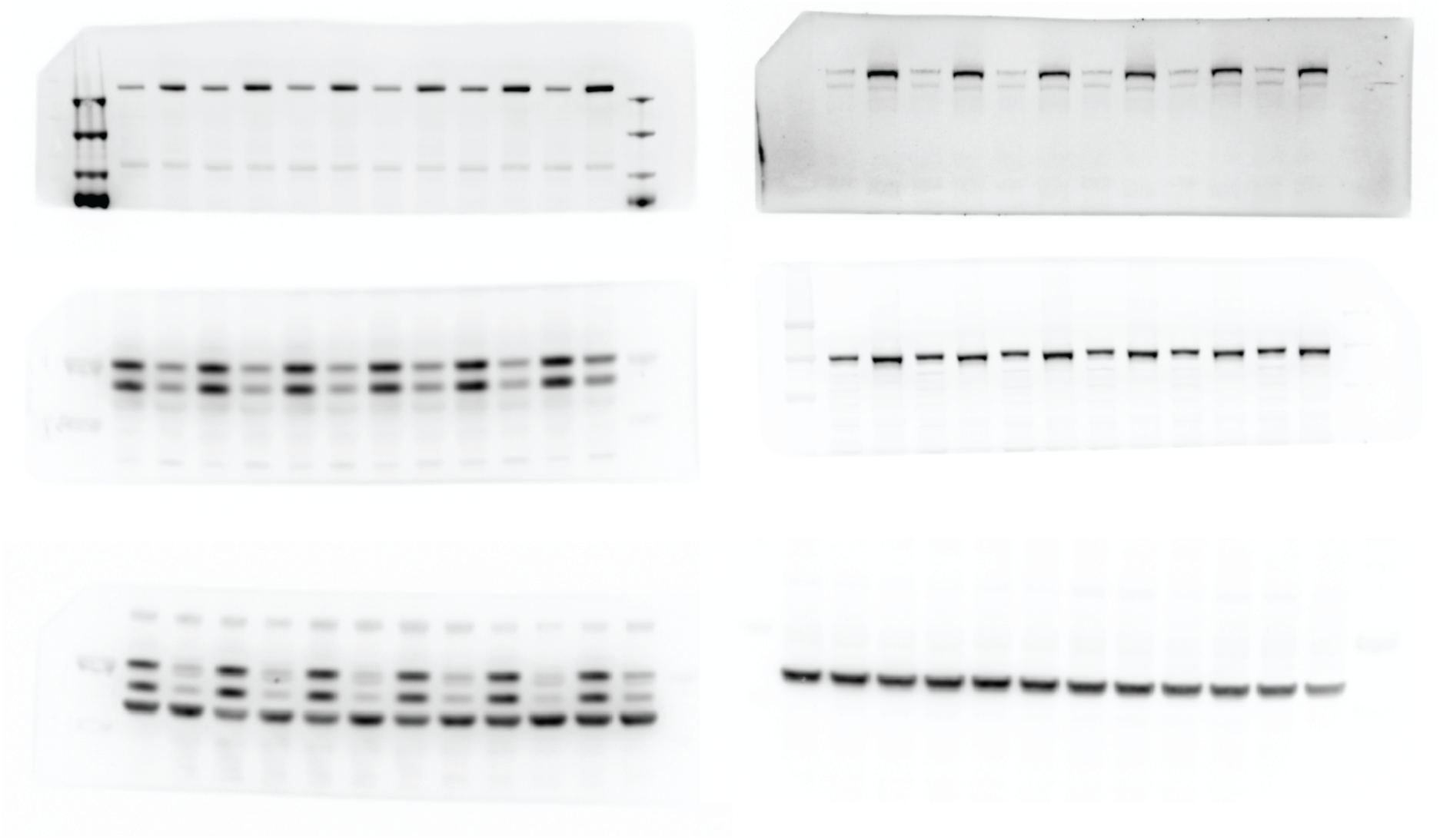
Uncropped western blots shown in figure 1.

## Reference

Aguzzi, A. and Y. Liu (2017). "A role for astroglia in prion diseases." J Exp Med 214(12): 3477–3479.

Akhtar, S., A. Wenborn, S. Brandner, J. Collinge and S. E. Lloyd (2011). "Sex effects in mouse prion disease incubation time." PLoS One 6(12): e28741.

Alonso, G. (2005). "NG2 proteoglycan-expressing cells of the adult rat brain: possible involvement in the formation of glial scar astrocytes following stab wound." Glia 49(3): 318–338.

Avar, M., D. Heinzer, N. Steinke, B. Dogancay, R. Moos, S. Lugan, C. Cosenza, S. Hornemann, O. Andreoletti and A. Aguzzi (2020). "Prion infection, transmission, and cytopathology modeled in a low-biohazard human cell line." Life Sci Alliance 3(8).

Bonfill-Teixidor, E., A. Otxoa-de-Amezaga, M. Font-Nieves, M. G. Sans-Fons and A. M. Planas (2017). "Differential expression of E-type prostanoid receptors 2 and 4 in microglia stimulated with lipopolysaccharide." J Neuroinflammation 14(1): 3.

Bradford, B. M., L. I. McGuire, D. A. Hume, C. Pridans and N. A. Mabbott (2022). "Microglia deficiency accelerates prion disease but does not enhance prion accumulation in the brain." Glia 70(11): 2169–2187.

Brandner, S., S. Isenmann, A. Raeber, M. Fischer, A. Sailer, Y. Kobayashi, S. Marino, C. Weissmann and A. Aguzzi (1996). "Normal host prion protein necessary for scrapie-induced neurotoxicity." Nature 379(6563): 339–343.

Brandner, S. and Z. Jaunmuktane (2017). "Prion disease: experimental models and reality." Acta Neuropathol 133(2): 197–222.

Bueler, H., A. Aguzzi, A. Sailer, R. A. Greiner, P. Autenried, M. Aguet and C. Weissmann (1993). "Mice devoid of PrP are resistant to scrapie." Cell 73(7): 1339–1347.

Calver, A. R., A. C. Hall, W. P. Yu, F. S. Walsh, J. K. Heath, C. Betsholtz and W. D. Richardson (1998). "Oligodendrocyte population dynamics and the role of PDGF in vivo." Neuron 20(5): 869–882.

Carroll, J. A., B. Race, K. Williams, J. Striebel and B. Chesebro (2018). "Microglia Are Critical in Host Defense against Prion Disease." J Virol 92(15).

Chen, W. T., A. Lu, K. Craessaerts, B. Pavie, C. Sala Frigerio, N. Corthout, X. Qian, J. Lalakova, M. Kuhnemund, I. Voytyuk, L. Wolfs, R. Mancuso, E. Salta, S. Balusu, A. Snellinx, S. Munck, A. Jurek, J. Fernandez Navarro, T. C. Saido, I. Huitinga, J. Lundeberg, M. Fiers and B. De Strooper (2020). "Spatial Transcriptomics and In Situ Sequencing to Study Alzheimer’s Disease." Cell 182(4): 976–991 e919.

Dimitriadis, A., F. Zhang, T. Murphy, T. Trainer, Z. Jaunmuktane, C. Schmidt, T. Nazari, J. Linehan, S. Brandner, J. Collinge, S. Mead and E. Viré (2022). "Single-nuclei transcriptomics of mammalian prion diseases identifies dynamic gene signatures shared between species." bioRxiv: 2022.2009.2013.507650.

Ettle, B., J. C. M. Schlachetzki and J. Winkler (2016). "Oligodendroglia and Myelin in Neurodegenerative Diseases: More Than Just Bystanders?" Mol Neurobiol 53(5): 3046–3062.

Falsig, J. and A. Aguzzi (2008). "The prion organotypic slice culture assay--POSCA." Nat Protoc 3(4): 555-562.

Fischer, M., T. Rulicke, A. Raeber, A. Sailer, M. Moser, B. Oesch, S. Brandner, A. Aguzzi and C. Weissmann (1996). "Prion protein (PrP) with amino-proximal deletions restoring susceptibility of PrP knockout mice to scrapie." EMBO J 15(6): 1255–1264.

Fruhbeis, C., W. P. Kuo-Elsner, C. Muller, K. Barth, L. Peris, S. Tenzer, W. Mobius, H. B. Werner, K. A. Nave, D. Frohlich and E. M. Kramer-Albers (2020). "Oligodendrocytes support axonal transport and maintenance via exosome secretion." PLoS Biol 18(12): e3000621.

Fuhrmann, M., G. Mitteregger, H. Kretzschmar and J. Herms (2007). "Dendritic pathology in prion disease starts at the synaptic spine." J Neurosci 27(23): 6224–6233.

Funfschilling, U., L. M. Supplie, D. Mahad, S. Boretius, A. S. Saab, J. Edgar, B. G. Brinkmann, C. M. Kassmann, I. D. Tzvetanova, W. Mobius, F. Diaz, D. Meijer, U. Suter, B. Hamprecht, M. W. Sereda, C. T. Moraes, J. Frahm, S. Goebbels and K. A. Nave (2012). "Glycolytic oligodendrocytes maintain myelin and long-term axonal integrity." Nature 485(7399): 517–521.

Jin, S., C. F. Guerrero-Juarez, L. Zhang, I. Chang, R. Ramos, C. H. Kuan, P. Myung, M. V. Plikus and Q. Nie (2021). "Inference and analysis of cell-cell communication using CellChat." Nat Commun 12(1): 1088.

Jin, X., T. R. Riew, H. L. Kim, J. H. Choi and M. Y. Lee (2018). "Morphological characterization of NG2 glia and their association with neuroglial cells in the 3-nitropropionic acid-lesioned striatum of rat." Sci Rep 8(1): 5942.

Johansson, J. U., S. Pradhan, L. A. Lokteva, N. S. Woodling, N. Ko, H. D. Brown, Q. Wang, C. Loh, E. Cekanaviciute, M. Buckwalter, A. B. Manning-Bog and K. I. Andreasson (2013). "Suppression of inflammation with conditional deletion of the prostaglandin E2 EP2 receptor in macrophages and brain microglia." J Neurosci 33(40): 16016–16032.

Keren-Shaul, H., A. Spinrad, A. Weiner, O. Matcovitch-Natan, R. Dvir-Szternfeld, T. K. Ulland, E. David, K. Baruch, D. Lara-Astaiso, B. Toth, S. Itzkovitz, M. Colonna, M. Schwartz and I. Amit (2017). "A Unique Microglia Type Associated with Restricting Development of Alzheimer’s Disease." Cell 169(7): 1276–1290 e1217.

Korsunsky, I., N. Millard, J. Fan, K. Slowikowski, F. Zhang, K. Wei, Y. Baglaenko, M. Brenner, P. R. Loh and S. Raychaudhuri (2019). "Fast, sensitive and accurate integration of single-cell data with Harmony." Nat Methods 16(12): 1289–1296.

Larsson, K., J. Steinmetz, F. Bergqvist, S. Arefin, L. Spahiu, J. Wannberg, S. C. Pawelzik, R. Morgenstern, P. Stenberg, K. Kublickiene, M. Korotkova and P. J. Jakobsson (2019). "Biological characterization of new inhibitors of microsomal PGE synthase-1 in preclinical models of inflammation and vascular tone." Br J Pharmacol 176(24): 4625–4638.

Lee, Y., B. M. Morrison, Y. Li, S. Lengacher, M. H. Farah, P. N. Hoffman, Y. Liu, A. Tsingalia, L. Jin, P. W. Zhang, L. Pellerin, P. J. Magistretti and J. D. Rothstein (2012). "Oligodendroglia metabolically support axons and contribute to neurodegeneration." Nature 487(7408): 443–448.

Liu, Y. and A. Aguzzi (2019). "Immunotherapy for neurodegeneration?" Science 364(6436): 130–131.

Liu, Y. and A. Aguzzi (2020). "NG2 glia are required for maintaining microglia homeostatic state." Glia 68(2): 345–355.

Liu, Y., A. Senatore, S. Sorce, M. Nuvolone, J. Guo, Z. H. Gumus and A. Aguzzi (2022). "Brain aging is faithfully modelled in organotypic brain slices and accelerated by prions." Commun Biol 5(1): 557.

Liu, Y., S. Sorce, M. Nuvolone, J. Domange and A. Aguzzi (2018). "Lymphocyte activation gene 3 (Lag3) expression is increased in prion infections but does not modify disease progression." Sci Rep 8(1): 14600.

Liu, Y. and J. Zhou (2013). "Oligodendrocytes in neurodegenerative diseases." Frontiers in Biology 8(2): 127–133.

Loeuillet, C., P. Y. Boelle, C. Lemaire-Vieille, M. Baldazza, P. Naquet, P. Chambon, M. F. Cesbron-Delauw, A. J. Valleron, J. Gagnon and J. Y. Cesbron (2010). "Sex effect in mouse and human prion disease." J Infect Dis 202(4): 648–654.

Mathys, H., C. Adaikkan, F. Gao, J. Z. Young, E. Manet, M. Hemberg, P. L. De Jager, R. M. Ransohoff, A. Regev and L. H. Tsai (2017). "Temporal Tracking of Microglia Activation in Neurodegeneration at Single-Cell Resolution." Cell Rep 21(2): 366–380.

Mathys, H., J. Davila-Velderrain, Z. Peng, F. Gao, S. Mohammadi, J. Z. Young, M. Menon, L. He, F. Abdurrob, X. Jiang, A. J. Martorell, R. M. Ransohoff, B. P. Hafler, D. A. Bennett, M. Kellis and L. H. Tsai (2019). "Single-cell transcriptomic analysis of Alzheimer’s disease." Nature 570(7761): 332–337.

Minghetti, L., F. Cardone, A. Greco, M. Puopolo, G. Levi, A. J. Green, R. Knight and M. Pocchiari (2002). "Increased CSF levels of prostaglandin E(2) in variant Creutzfeldt-Jakob disease." Neurology 58(1): 127–129.

Minghetti, L., A. Greco, F. Cardone, M. Puopolo, A. Ladogana, S. Almonti, C. Cunningham, V. H. Perry, M. Pocchiari and G. Levi (2000). "Increased brain synthesis of prostaglandin E2 and F2-isoprostane in human and experimental transmissible spongiform encephalopathies." J Neuropathol Exp Neurol 59(10): 866–871.

Minikel, E. V., H. T. Zhao, J. Le, J. O’Moore, R. Pitstick, S. Graffam, G. A. Carlson, M. P. Kavanaugh, J. Kriz, J. B. Kim, J. Ma, H. Wille, J. Aiken, D. McKenzie, K. Doh-Ura, M. Beck, R. O’Keefe, J. Stathopoulos, T. Caron, S. L. Schreiber, J. B. Carroll, H. B. Kordasiewicz, D. E. Cabin and S. M. Vallabh (2020). "Prion protein lowering is a disease-modifying therapy across prion disease stages, strains and endpoints." Nucleic Acids Res 48(19): 10615-10631.

Nakano, M., Y. Tamura, M. Yamato, S. Kume, A. Eguchi, K. Takata, Y. Watanabe and Y. Kataoka (2017). "NG2 glial cells regulate neuroimmunological responses to maintain neuronal function and survival." Sci Rep 7: 42041.

Pandey, S., K. Shen, S. H. Lee, Y. A. Shen, Y. Wang, M. Otero-Garcia, N. Kotova, S. T. Vito, B. I. Laufer, D. F. Newton, M. G. Rezzonico, J. E. Hanson, J. S. Kaminker, C. J. Bohlen, T. J. Yuen and B. A. Friedman (2022). "Disease-associated oligodendrocyte responses across neurodegenerative diseases." Cell Rep 40(8): 111189.

Slota, J. A., B. V. Sajesh, K. F. Frost, S. J. Medina and S. A. Booth (2022). "Dysregulation of neuroprotective astrocytes, a spectrum of microglial activation states, and altered hippocampal neurogenesis are revealed by single-cell RNA sequencing in prion disease." Acta Neuropathol Commun 10(1): 161.

Sorce, S., M. Nuvolone, G. Russo, A. Chincisan, D. Heinzer, M. Avar, M. Pfammatter, P. Schwarz, M. Delic, M. Muller, S. Hornemann, D. Sanoudou, C. Scheckel and A. Aguzzi (2020). "Genome-wide transcriptomics identifies an early preclinical signature of prion infection." PLoS Pathog 16(6): e1008653.

Stuart, T., A. Butler, P. Hoffman, C. Hafemeister, E. Papalexi, W. M. Mauck, 3rd, Y. Hao, M. Stoeckius, P. Smibert and R. Satija (2019). "Comprehensive Integration of Single-Cell Data." Cell 177(7): 1888-1902 e1821.

Wareham, L. K., S. A. Liddelow, S. Temple, L. I. Benowitz, A. Di Polo, C. Wellington, J. L. Goldberg, Z. He, X. Duan, G. Bu, A. A. Davis, K. Shekhar, A. Torre, D. C. Chan, M. V. Canto-Soler, J. G. Flanagan, P. Subramanian, S. Rossi, T. Brunner, D. E. Bovenkamp and D. J. Calkins (2022). "Solving neurodegeneration: common mechanisms and strategies for new treatments." Mol Neurodegener 17(1): 23.

Wolock, S. L., R. Lopez and A. M. Klein (2019). "Scrublet: Computational Identification of Cell Doublets in Single-Cell Transcriptomic Data." Cell Syst 8(4): 281–291 e289.

Woodling, N. S., Q. Wang, P. G. Priyam, P. Larkin, J. Shi, J. U. Johansson, I. Zagol-Ikapitte, O. Boutaud and K. I. Andreasson (2014). "Suppression of Alzheimer-associated inflammation by microglial prostaglandin-E2 EP4 receptor signaling." J Neurosci 34(17): 5882–5894.

Yang, S., S. E. Corbett, Y. Koga, Z. Wang, W. E. Johnson, M. Yajima and J. D. Campbell (2020). "Decontamination of ambient RNA in single-cell RNA-seq with DecontX." Genome Biol 21(1): 57.

Zhu, C., U. S. Herrmann, J. Falsig, I. Abakumova, M. Nuvolone, P. Schwarz, K. Frauenknecht, E. J. Rushing and A. Aguzzi (2016). "A neuroprotective role for microglia in prion diseases." J Exp Med 213(6): 1047–1059.

